# Live cell imaging of ATP levels reveals metabolic compartmentalization within motoneurons and early metabolic changes in *FUS* ALS motoneurons

**DOI:** 10.1101/2023.03.22.533787

**Authors:** Vitaly Zimyanin, Anne-Marie Pielka, Hannes Glaß, Julia Japtok, Melanie Martin, Andreas Deussen, Barbara Szewczyk, Chris Deppmann, Eli Zunder, Peter M. Andersen, Tobias M. Boeckers, Jared Sterneckert, Stefanie Redemann, Alexander Storch, Andreas Hermann

**Affiliations:** Department of Molecular Physiology and Biological Physics, University of Virginia, School of Medicine, Charlottesville, VA, USA; Center for Membrane and Cell Physiology, University of Virginia, School of Medicine, Charlottesville, VA, USA; Department of Neurology, Technische Universität Dresden, Dresden, Germany; Translational Neurodegeneration Section „Albrecht Kossel“, Department of Neurology, University Medical Center Rostock, University of Rostock, Rostock, Germany; Institute of Physiology, Technische Universität Dresden, Germany; Department of Biology, Graduate School of Arts and Sciences, University of Virginia, Charlottesville, VA, USA; Department of Biomedical Engineering, University of Virginia, School of Medicine, Charlottesville, VA, USA; Deutsches Zentrum für Neurodegenerative Erkrankungen (DZNE), Ulm Site, Ulm, Germany; Centre for Regenerative Therapies, Technische Universita dresden.de Dresden, Dresden, Germany; Department of Cell Biology, University of Virginia, School of Medicine, Charlottesville, VA, USA; Deutsches Zentrum für Neurodegenerative Erkrankungen (DZNE) Rostock/Greifswald, 18147 Rostock, Germany; Department of Clinical Sciences, Neurosciences, Umeå University,SE-901 85 Umeå, Sweden; Center for Transdisciplinary Neurosciences Rostock (CTNR), University Medical Center Rostock, University of Rostock, 18147 Rostock, Germany; Department of Neurology, University of Rostock, Rostock, Germany; Institute for Anatomy and Cell Biology, Ulm University, Ulm, Germany

**Keywords:** Amyotrophic lateral sclerosis, Mitochondria, Metabolism

## Abstract

Motoneurons are one of the highest energy demanding cell types and a primary target in Amyotrophic lateral sclerosis (ALS), a debilitating and lethal neurodegenerative disorder without currently available effective treatments. Disruption of mitochondrial ultra-structure, transport and metabolism is a commonly reported phenotype in ALS models and can critically affect survival and proper function of motor neurons. However, how changes in metabolic rates contribute to ALS progression are not fully understood yet. Here we utilize hiPCS derived motoneuron cultures and live imaging quantitative techniques to evaluate metabolic rates in Fused in Sarcoma (FUS)-ALS model cells. We show that differentiation and maturation of motoneurons is accompanied by an overall upregulation of mitochondrial components and significant increase in metabolic rates that corresponds to their high energy-demanding state. Detailed compartment-specific live measurements using a fluorescent ATP sensor and FLIM imaging show significantly lower levels of ATP in the somas of cells carrying FUS-ALS mutations. These changes lead to the increased vulnerability of disease motoneurons to further metabolic challenges with mitochondrial inhibitors and could be due to the disruption of mitochondrial inner membrane integrity and an increase in its proton leakage. Furthermore, our measurements demonstrate heterogeneity between axonal and somatic compartments with lower relative levels of ATP in axons. Our observations strongly support the hypothesis that mutated FUS impacts metabolic states of motoneurons and makes them more susceptible to further neurodegenerative mechanisms.

## 1. Introduction

Amyotrophic lateral sclerosis (ALS) is a severe and still incurable disease that is characterized by a more or less selective degeneration of upper and lower motor neurons (MN) in the motor cortex, brainstem and spinal cord. Degeneration of motoneurons leads to denervation of muscles and to death within two to five years after symptom onset [1]. The majority of ALS cases are sporadic with only 10% of them being monogenetically inherited. Genes for ‘SUPEROXIDE DISMUTASE 1’ (*SOD1*), ‘FUSED IN SARCOMA’ (*FUS*), ‘TAR DNA BINDING PROTEIN’ (*TARDBP*) or a hexanucleotide repeat expansion in the ‘CHROMOSOME 9 OPEN READING FRAME 72’ (*C9ORF7*2) gene are the source of the majority of ALS heritable cases [2-4]. Modelling of genetic ALS cases using various *in vivo* and *in vitro* systems has provided an invaluable insight into molecular mechanisms of the disease and at the same time highlights the challenges that are inherent to our understanding and treatment of most neurodegenerative diseases. These are: complexity and broad scope of underlying molecular changes, long-term and pre-symptomatic disruption of cell homeostasis that becomes critical with age and/or extra stressors, yet a remarkable cell type and region specificity of neuron type vulnerable to degeneration.

Fast-fatigable motoneurons, which have highest energy demands, get affected first and are most vulnerable in ALS, while slow firing motoneurons and sensory neurons remain relatively intact [5-7]. It has been therefore proposed that bioenergetic failure is one of the critical factors behind MN degeneration [8-12]. In support of this hypothesis, abnormal morphologies of mitochondria and disrupted function have been reported from postmortem samples of ALS patients and are an early marker in a number of rodent ALS models [6, 11, 13-16] and in iPSC-derived motoneurons [10, 17-20]. Changes in global metabolism and ability to catabolize glucose by MN correlate with the severity of ALS progression [21, 22]. ALS-causing mutations in *FUS* were recently reported to disrupt several energy demanding processes in iPSC-derived MNs such as neuronal firing rates and axonal transport [20, 23, 24]. Finally, FUS overexpression or mutations in various model systems disrupt mitochondrial structure and bioenergetics [13-16, 25].

The exact role of FUS in regulating metabolism is far from clear. The *FUS* gene encodes a primarily nuclear protein that contains DNA/RNA binding- and low-complexity-domains and plays a role in regulating transcription, DNA damage, RNA splicing, nucleocytoplasmic trafficking, ER stress, protein translation and nonsense mediated decay [26-30]. FUS binding was mapped to multiple sites on chromosomes and more than a thousand long coding mRNAs, where it was shown to regulate their splicing [31-34]. Thus, it can directly or indirectly affect multiple components involved in metabolism and its regulation. Recent examples suggest a specific enrichment of mitochondrial respiratory chain complexes transcripts in FUS aggregates resulting in downregulation of their protein level [11]. In addition, FUS has been shown to play a very direct role in regulation of metabolism. FUS mis-localization from the nucleus to cytoplasm due to mutations in nuclear-localization signal sequence or FUS overexpression results in an increased interaction of FUS with enzymes involved in glucose metabolism or binding of FUS to ATP synthase beta subunit and induction of mitochondrial unfolded protein response in cellular and animal models [14, 15].

Metabolism rates in turn affect the function of FUS itself. FUS protein Low-Complexity domains (LCDs) are essential for its ability to undergo liquid-liquid phase separation, a mechanism that is broadly utilized in cells to concentrate certain molecules or functional complexes in so called membraneless organelles [35-38]. ALS-specific mutations in FUS that change its protein-protein interactions properties or cause mis-localization to the cytoplasm and therefore increase in protein’s local concentration were proposed to trigger changes in phase-transition properties of high-order protein complexes. These changes were shown to cause maturation of FUS first into hydrogel like state and later into solid-like irreversible fibrillar aggregates, in a so-called liquid-to-solid transition [37, 39-43]. Such protein aggregation is a principle hallmark of ALS and is characteristic not only for FUS and ALS but in fact for most neurodegenerative diseases. The metabolic state of the cell plays a critical role in controlling such aggregate formation. Chaperone and proteasome machineries bind and neutralize misfolded proteins via refolding or degradation functions at a high energy cost to the cell [44-49]. Furthermore, ATP levels in the cytoplasm have been previously shown to regulate the state of cytoplasm and stress granules under stress conditions [50, 51]. Very recent *in vitro* studies with ND LCDs like FUS strongly suggest that the global metabolic state could be important in preventing aberrant assemblies of LCD-s via hydrotrope properties of ATP [38].

Overall, metabolism can be an essential factor contributing to ALS progression with multiple and multilayered input from mis-regulation of ALS-causing gene targets and age-related changes in metabolism and cell homeostasis. Surprisingly, despite all of the reported phenotypes at least one recent publication was not able to link individual FUS mutations to global changes in MN metabolism [10]. This might be due to the analysis of mixed cultures and thus measurements might be masked by non motoneuronal cells. It is possible however that without FUS overexpression and the additional stress often utilized in approaches to model FUS ALS, effects of FUS mutations could be small or only occur in sub-compartments of MN, e.g. the axon. The latter would indeed fit well to the mainly distal axonal trafficking deficits of mitochondria and distal axonal mitochondrial depolarization found in FUS iPSC-derive MN [23].

Here we utilized human-induced pluripotent stem cell (iPSC)-derived MNs and used a range of techniques to re-visit measurements of metabolic rates in motoneurons. First, we characterized potential metabolic changes that occur during MN differentiation from neural precursor cells (NPC) to fully mature motoneurons and if these changes were affected by FUS mutations. We then used *in vivo* imaging of ATP-sensors to measure individual motoneurons as well as to increase the sensitivity of our measurements and assess relative ATP concentration changes in specific compartments of neurons: soma and distal axons. We showed that differentiation and maturation of motoneurons was accompanied by a global upregulation of metabolism and many of the tested mitochondrial components. We detected clear differences between somatic and distal axonal compartments in all motoneurons and showed early significant changes in ATP levels in the soma of motoneurons carrying FUS-ALS mutants. This early difference were sufficient to affect MN survival upon additional metabolic stress and occured most likely due to additional proton leakage in mitochondrial inner membrane. Furthermore, long-term culturing of motoneurons led to a significant drop in ATP concentration across all of the cell compartments. We propose that such changes can significantly exacerbate ALS-related disruption of cell-homeostasis and would be consistent with general etiology of most neurodegenerative diserases (NDs), which often start as axonopathies and are assumed to be sensitive to global slowing of metabolism and other age-related compounding factors.

## 2. Materials and Methods

### 2.1. Cell lines

All cell lines were obtained from skin biopsies of patients or healthy volunteers, or generated using CRISP/Cas mutagenesis and have been described before (Table 1). The performed procedures were in accordance with the Declaration of Helsinki (WMA, 1964) and approved by the Ethical Committee of the Technische Universität Dresden, Germany (EK 393122012 and EK 45022009) and by the Swedish Ethical Committee for Medical Research (Nr 94-135 to perform genetic research including for FUS and Nr 2018-494-32M to perform *in vitro* lab studies on cell lines derived from patients with ALS). Written informed consent was obtained from all participants including for publication of any research results. All fibroblast lines were checked for Mycoplasma sp. prior to and after reprogramming and afterwards, routine checks for Mycoplasma were done every three to six months. We used the Mycoplasma Detection kit according to manufacturer’s instructions (Firma Venor GeM, No 11–1025).

**Table 1.** Cell lines used in this study.

|  | FUS Line | Sex | Age at Biopsy | Mutation | Primarily Characterized in |
| --- | --- | --- | --- | --- | --- |
| Wt | Ctrl1 | Female | 43 | - | [52] |
| Wt | Ctrl2 | Female | 49 | - | [52] |
| Wt | Ctrl3 | Female | 48 | - | [65] |
| Wt | Ctrl4-GFP | - | - | - | [23] |
| Mt | FUS-1 | Female | 58 | R521C | [65] |
| Mt | FUS-2 | Female | 65 | R521L | [23] |
| Mt | FUS-3-GFP | - | - | P525L | [23] |
| Mt | FUS-4 | Male | 29 | R495QfsX527 | [23, 65] |
| Mt | FUS-5 | Male |  | Q23L | This study |

### 2.2. Fibroblasts reprogramming

Patient fibroblasts were reprogrammed using pMX-based retroviral vectors encoding the human cDNAs of OCT4, SOX2, KLF4 and cMYC (pMX vectors). Vectors were co-transfected with packaging-defective helper plasmids into 293T cells using Fugene 6 transfection reagent (Roche). Fibroblasts were plated at a density of 50,000 cells/well on 0.1% gelatin-coated 6-well plates and infected three times with a viral cocktail containing vectors expressing OCT4:SOX2:KLF4:cMYC in a 2:1:1:1 ratio in the presence of 6 µg/ml protamine sulfate (Sigma Aldrich) and 5 ng/ml FGF2 (Peprotech). Infected fibroblasts were plated onto matrigel coated plates at a density of 10,000 cells per 1w/6w in fibroblast media containing 1 mM VPA an incubated for 48h. Afterwards, medium was changed to TeSR-E7 (Stemcell Technologies) containing 1 mM VPA. Media was changed every day to the same conditions. VPA was added 7-10 days. Afterwards cells were cultured in TeSR-E7 only. iPSC-like clusters started to appear at day 7 post infection, were manually picked 14 days post-infection and plated onto matrigel in regular TeSR-E8 (stemcell Technologies) medium in addition of 10 µM Y-27632 (Ascent Scientific). Stable clones were routinely passaged onto matrigel using 1 mg/ml collagenase type IV (Invitrogen) and addition of 10 µM Y-27632 (Ascent Scientific) for the first 48 h after passaging. Media change with addition of fresh FGF2 was performed every day.

### 2.3. NPCs and motoneuron generation

The generation of human NPCs and MNs was accomplished following the modifications protocol from [52] and described also in [23]: In brief, colonies of iPSCs were collected and stem cell medium, containing 10 µM SB-431542, 1 µM Dorsomorphin, 3 µM CHIR 99021 and 0.5 µM pumorphamine (PMA), was added. After 2 days hESC medium was replaced with N2B27 consisting of the aforementioned factors and DMEM-F12/Neurobasal 50:50 with 1:200 N_2_ Supplement, 1:100 B27 lacking Vitamin A and 1% penicillin / streptomycin / glutamine. On day 4 150 µM ascorbic acid was added while Dorsomorphin and SB-431542 were withdrawn. 2 Days later the EBs were mechanically separated and replated on Matrigel coated dishes. For this purpose Matrigel was diluted (1:100) in DMEM-F12 and kept on the dishes over night at room temperature. Possessing a ventralized and caudalized character the arising so called small molecule NPCs (smNPC) formed homogenous colonies during the course of further cultivation. It was necessary to split them at a ratio of 1:10–1:20 once a week using Accutase for 10 min at 37° C. Final MN differentiation was induced by treatment with 1 µM PMA in N2B27 exclusively. After 2 days 1 µM retinoic acid (RA) was added. On day 9 another split step was performed to seed them on a desired cell culture system. Furthermore the medium was modified to induce neural maturation. For this purpose the developing neurons were treated with N2B27 containing 10 ng/µl BDNF, 500 µM dbcAMP and 10 ng/µl GDNF. Following this protocol it was possible to keep the cells in culture for up to 15-20 weeks.

### 2.4. Growth in motoneurons in microfluidic chambers

The MFCs were purchased from Xona (RD900). At first, Nunc glass bottom dishes with an inner diameter of 27 mm were coated with Poly-L-Ornithine (Sigma-Aldrich P4957, 0.01% stock diluted 1:3 in PBS) overnight at 37° C. After 3 steps of washing with sterile water, they were kept under the sterile hood for air drying. MFCs were sterilized with 70% Ethanol and also left drying. Next, the MFCs were dropped onto the dishes and carefully pressed on the glass surface for firm adherence. The system was then perfused with Laminin (Roche 11243217001, 0.5 mg/ml stock diluted 1:50 in PBS) for 3 h at 37° C. For seeding cells, the system was once washed with medium and then 10 µl containing a high concentration 3×10^5^ cells (3×10^7^ cells/ml) were directly injected into the main channel connecting two wells. After allowing for cell attachment over 30–60 min in the incubator, the still empty wells were filled up with maturation medium. This method had the advantage of increasing the density of neurons in direct juxta-position to microchannel entries whereas the wells remained cell-free, thereby reducing the medium turnover to a minimum. To avoid drying out, PBS was added around the MFCs. Two days after seeding, the medium was replaced in a manner which gave the neurons a guidance cue for growing through the microchannels. Specifically, a growth factor gradient was established by adding 100 µl N2B27 with 500 µM dbcAMP to the proximal seeding site and 200 µl N2B27 with 500 µM dbcAMP, 10 ng/µl BNDF, 10 ng/µl GDNF and 100 ng/µl NGF to the distal exit site. The medium was replaced in this manner every third day. After 7 days, the first axons began spreading out at the exit site and cells were typically maintained for up to six weeks. Mitochondrial inhibitors Oligomycin A 10 µM (10 mM stock) and CCCP 10 uM (10 mM stock) were added directly into the media 1 hour prior to imaging.

### 2.5. High performance liquid chromatography

Native AMP, ADP and ATP were derivatized to 1,N6-etheno derivatives of endogenous adenine nucleotides as described before [53] with some modifications. 143 µl of the diluted samples (1:2 with bidistilled. water) were incubated with 51 µl of citrate phosphate buffer (62 and 76 mM, respectively; pH 4.0) and 6 µl chloracetaldehyde at 80° C for 40 min. Derivatized samples were immediately cooled to 4° C and stored at -20° C until further analysis with HPLC. High performance liquid chromatography (HPLC) for quantification of the nucleotides was performed on a Waters Alliance 2690 HPLC (Waters, Milford, Mass., USA) coupled to a Waters 2475 fluorescence detector (wavelength ex=280 nm; wavelength em=410 nm) as described previously [53] with some modifications. In brief, separations were carried out on a Waters XTerra MS C18, 4.6×50 mm I.D. column with a particle size of 5 µm. A guard column packed with the same sorbent was used. 10 µl of sample was injected by an autosampler and compounds were eluted with a flow rate of 0.75 ml/min using a binary tetrabutylammoniumhydrogensulfate (TBAS)/acetonitrile gradient. Eluent A contained 6% acetonitrile in TBAS buffer and eluent B contained 65% acetonitrile in TBAS buffer. Initial conditions were 100% A, changed to 66% A within 5.6 min, maintained for 2.4 min, then 100% A was reestablished within 0.5 min and the column was equilibrated for 6.5 min. External standards of known concentrations were used to check retention times and to permit sample quantification based on the analysis of peak area. Empower2 (Waters) was used for report. Тhe ATP/ADP ratio and adenylate energy charge (AEC) has been adopted to describe the energetic status of a cell or tissue preparation. The AEC is the ratio of the compete adenylate pool defined as AEC=[[ATP+(0.5ADP)]/(AMP+ADP+ATP)] [54-56].

### 2.6. FLIM measurements

For fluorescence lifetime imaging (FLIM) imaging we used inverted Laser Scanning Confocal microscope LSM 780/FLIM inverse Zeiss microscope in the imaging facility of BIOTEC in Dresden with a temperature and CO2 controlled chamber. Microscope was equipped with Becker & Hickl dual channel FLIM unit and uses 440 nm pulsed Laser Diode. For A-team imaging we use used CFP/YFP double cube set F46-001 and 40x/1.2 LD LCI Plan-Apochromat lense and inverted Laser Scanning Confocal microscope. FLIM Data fitting is based on the Becker & Hickl handbook and SPCImage software (v. 5.5, Becker & Hickl) software. We used two components incomplete model to fit CFP and YFP FLIM images. The offset and scattering are set to 0. Shift is optimized to make sure the Chi2 as close as to 1 (between 0.7 and 2).

### 2.7. FLIM Processing and Analysis

FLIM processing follows the previous published papers [57, 58] with an important advance in normalization of photon reference images to compensate for varying intensities (FIJI, Plugins -> Integral Image Filters -> Normalize Local Contrast): followed by zeroing the nucleus; cell segmentation and creating single pixel ROIs by a ImageJ/FIJI custom plugin (Suppl. Fig. 5). The purpose of this sequence is to create pixel locations by X-Y coordinates, specific for cytosolic cell body or axonal MN and exclude background and noise measurements. Those locations are then applied to the FLIM data to extract any of the FLIM parameters in the data pool. A number of parameters were generated including photon images, t_1_, t_2_, a_1_%, a_2_%, and χ^2^ for each pixel of each channel and T_m_ was calculated as T_m_=a_1_%×t_1_+a_2_%×t_2._ A custom made application Flimanalyzer (https://flimanalyzer.readthedocs.io/en/latest/) based on Python code and a custom made in UVA KECK centre ultimately analyzes different data combinations to produce ratios, means, medians, which are further charted either with Flimanalyzer or PRIZM as bar graphs, frequency histograms or normalized kernel density estimate (KDE) plots. We used a T-test ‘Two-sample assuming unequal variances’ or two way ANOVA for significance tests with the significance cut off level of <5% (*p*<0.05).

### 2.8. Immunoblotting

For mitochondrial OXPHOS protein expression, proteins were extracted in RIPA-buffer and protein concentration determined with BCA assay (Pierce). Cell lysates were then diluted in SDS-PAGE loading buffer (Roti-Load, K929.1, Carl Roth), and SDS-PAGE was performed on 20 µg of cell lysate per sample on Tris-glycine-SDS polyacrylamide gels (4561095, Biorad) then transferred to PVDF (1620177, Biorad) using standard techniques. Primary antibodies used were against the mitochondrial respiratory chain subunits (mitochondrial OXPHOS antibody cocktail containing cytochrome c oxidase subunit 2 [COX2], cytochrome b-c1 complex subunit 2 [UQCRC2], succinate dehydrogenase [ubiquinone] flavoprotein subunit B [SDHB], NADH dehydrogenase [ubiquinone] 1 beta subcomplex subunit 8 [NDUFB8] and ATP synthase subunit alpha [ATP5A]; Cat. #ab110411, Abcam). Total protein detection (REVERT Total Protein Stain Kit, P/N 926-11010;Li-Cor) was used to normalize for total cellular protein. Primary antibodies were detected with appropriate anti-mouse or anti-rabbit Infared IRDye 680 RD or 800 CW antibodies (Abcam). Protein band intensities were detected with the Odyssey 3.0 (Li-Cor Biosciences) and quantified using Image Lab 6.0 software (Biorad), and band intensity determined in the linear range was normalized to band intensity of total protein.

### 2.9. Isolation and analysis of RNA and DNA

For isolation of DNA and RNA, samples were purified on day 7 of expansion, day 7 of differentiation, and day 7,14,21 and 28 of maturation using AllPrep DNA/RNA/Protein (80004, Qiagen) according to the manufacturer’s instructions. To check the yield and purity of the RNA, the absorbance of the samples was measured in a measured in duplicate using Nanodrop ND 1000 (VWR International GmbH). For analysis by qPCR, RNA samples were converted to cDNA using the QuantiTect Reverse Transcription Kit (205313, Qiagen) into cDNA. For this purpose, the RNA samples were converted to cDNA depending on the absorbance results to a final concentration of 100 ng/µl, 300 ng/µl or 600 ng/µl were adjusted. In the thermal cycler, samples were incubated at 42 °C for 20 min, boiled at 95° C for 3 min and then stopped on ice and stored at -20° C. The qPCR was performed with the QuantiNova SYBR Green PCR Kit (208052, Qiagen). For this purpose 10 ng of each sample was used and applied in duplicate. The primer sequences and concentrations used are shown in the table. The qPCR started with a two-minute initiation step at 95° C. This was followed by denaturation for 5 s and 95° C and primer hybridization with extension of the target sequences for 10 s and 60° C. This cycle, without the initiation step, was repeated 38 times and a melting curve was generated at the end. The evaluation of the qPCR is done with the Rotor-Gene Q Series software. For each primer a concentration test was carried out for each primer before the standard curve was generated to determine the best possible threshold value through internal regression.

### 2.10. MTT assay

Cell viability status was measured with the colorimetric MTT assay kit (ab211091, Abcam), according to the manufacturer’s protocol. Briefly, motor neuron cells were seeded into 96-well plate (735-0465, VWR) at a density of 30×103 in 100 µL of medium per well in two replicates per each group treatment. After 07div or 28div, cells were incubated for 24 h with experimental factors (oligomycin A (75351, Sigma Aldrich); CCCP (carbonyl cyanide 3-chlorophenylhydrazone) (C2759-100MG, Sigma Aldrich); rotenone (MKBZ2534C, Sigma Aldrich); ddC (2’,3’-dideoxycytidine) (ab142240, Abcam); etoposide (Akos); 2-DG (Akos)), in standard conditions (at 37° C in a humidified atmosphere containing 5% CO2). After that time, the growth medium was removed from the wells and cells were incubated with equal volumes of growth medium and 3-(4,5-dimethylthiazol-2-yl)-2,5-diphenyltetrazolium bromide (MTT) reagent (50:50µl) for 3 h at 37° C. After incubation, formazan crystals were dissolved in 150 µl MTT solvent, and was wrapped with foil and shaken on an orbital shaker for 15 min. Absorbance at 570 nm was read on Microplate Reader (CM Sunrise™, Tecan) within one hour. Measurements within each line were normalized to the average values of mock treated wells of each line.

### 2.11. Seahorse measurement

Characteristic parameters of glycolysis and oxidative phosphorylation were assessed using Seahorse XFe97 Analyzer (Agilent). Motoneurons were generated according to 2.2. NPCs and Motoneuron generation. During the split on day 9 cells were transferred to a Seahorse XF96 cell culture plate, which was PLO/Laminin coated before. Prior to Seahorse measurement cells were washed for 30 min at 37° C with buffer free Seahorse XF base medium. Wells on the outside of the Seahorse XF96 cell culture plate were alternatingly filled with base medium and PBS. Injection ports were loaded to to block specific complexes. InjectionA - activation of glycolysis: 10 mM Glucose (Sigma)/ 1 mM Pyruvate (Thermo Fisher); InjectionB - Inhibition of Complex V: 1 µM Oligomycin A (Tocris); InjectionC – uncoupling: 0.7 µM FCCP (Tocris); Injection D – blocking of glycolysis and oxidatice phosphorylation: 50 mM 2-DG (Sigma)/ 1.5 µM Rotenone (Sigma)/ 2.5 µM Antimycin A (Sigma). Measurement consisted of five blocks: baseline, glycolysis activation, Blocking of complex V, Uncoupling, Blocking of glycolysis & oxidative phosphorylation. Each block was measured in triplicate. Each measurement consisted of 10 sec mixing, followed by 10 sec of settling and 3 min of measurement. In each experiment at least six wells per condition were measured in parallel and only wells were analyzed that showed normal run behavior (e.g. no sudden cell detachment). Extra cellular acidification rate (ECAR) and oxygen consumption rate (OCR) were calculated individually by robust regression for each well and measurement. Means were then calculated for biological replicates by pooling of the technical replicates of the same condition in each experiment.

### 2.12. NAD^+^/NADH ratio measurements

The experiment was performed according to the manufacturer’s description (NAD^+^/NADH Assay Kit, Abcam, ab176723). 5-7×10^6^ cells were scraped into PBS and centrifuged for 5 min at room temperature (1500 rpm). 5×10^6^ cells were then resuspended in 100 µl lysis buffer and incubated for 15 min at 37° C. Subsequently, the lysates were centrifuged at 1500 rpm for 5 min. The supernatant was then used for the NAD^+^/NADH assay. 25 µl NADH extraction buffer or NAD^+^ extraction buffer was added to each sample and then incubated for 15 min. For total NAD^+^/NADH measurement, samples were incubated with 25 µl NAD^+^/NADH control solution for 15min. Subsequently, the reaction was stopped with the opposite solution. 75 µl NADH reaction mixture was added to each well and the reaction was incubated for 30 min protected from light. Fluorescence was measured for up to 120 min on The Spark® multi-mode microplate reader (Tecan) at Ex/Em 540/590. Treatment with inhibitors were performed for 24 h with the following concentrations: PARG inhibitor Gallatonin (Santa Cruz, SC-202619) was used at 30 µM, PARP1 inhibitor ABT888 (Santa Cruz, SC-202901, 10 mg/ml in DMSO) was used at 2 µg/ml, FK866 (10 mM) that induces loss of NAD^+^ by inhibiting nicotinamide phosphoribosyltransferase was used at 10 µM.

### 2.13. Lentivirus production and delivery

A-team and A-team R122K/ R126K ORF from original constructs [59] were sub-cloned into pL lentiviral plasmid using Age1 and Sal1 restriction sites. Lentivirus was produced in HEK293T cells using 175 cm^2^ dishes. On the day before transfection, confluent HEK cells were trypsinized and seeded in IMDM complete media (+ 10% FBS) at 6-7 x 10^5^ cells/ml. IMDM (serum free) media and Poly(ethylenimine) (PEI) transfection reagent (Sigma) was used to transfect 10-12 µg of A-team control or A-team R122K/ R126K mutant plasmid together with pSPAX2 (6,5 µg) and pVSV-G (1-2 µg) plasmids for 30 mins. After washing of transfection media cells were then grown in complete IMDM (+10% FBS) media overnight. Next day media was replaced by serum-free complete Neurobasal medium with B-27 supplement and Pen/Strep (without glutamine). Virus was harvested at 24 h and 48 h time points, sterile filtered, aliquoted and stored at -80° C. Virus containing MN compatible media was added at a 1:10 ratio to freshly seeded neurons (DIV 0) and replaced with normal MN differentiation media after 24 h.

### 2.14. Statistics

Statistical analysis was done by either student t-test or one-way ANOVA followed by Bonferroni’s or Sidak method multiple comparisons test. Data were analyzed using the PRIZM, Excel or Flimanalyzer software. If not mentioned otherwise, all data are displayed as means±SEM. Significance level was set at *p*<0.05.

## 3. Results

### 3.1. FUS mutation has no effect on motoneuronal differentiation despite showing cytoplasmic FUS mislocalization

Recent protocols for the generation of patient specific motoneurons from iPSCs offer extremely powerful tools to model ALS in cell culture [60-64]. We previously generated human induced pluripotent stem cells (hiPSCs) either derived from fibroblasts of patients carrying ALS mutations or by CRISPR-based mutagenesis [23, 65, 66]. In this study we also have generated a new hiPSC FUS line from fibroblasts of a patient carrying a Q23L mutation in FUS/TLS. This newly derived hiPSCs were tested for the silencing of the transforming retrovirus, expression of standard pluripotency markers (Nanog, Oct4, Sox2 and Lin28A) and ability to successfully differentiate into and stain for for markers of all three germ layers (Suppl. Fig1). In our study we used the following hiPSCs with normal karyotype and confirmed mutations: *FUS*/ALS mutations in the nuclear-localization signal (NLS): P525L, R521C, R495QsfX527, as well as a mutation in a low complexity domain (LCD) of *FUS*: Q23L (Figure 1A-B, Table 1). We have previously shown that all NLS-mutation lines derive fully functional MNs and were tested for presence of phenotypes previously reported for ALS (Figure 1A) [23, 65, 67-69]. We could also confirm that MNs carrying mutations in the nuclear localization signal of FUS show its mis-localization to the cytoplasm (Figure 1C,D). To assess if changes in levels of FUS protein itself could be the cause of ALS-related cellular phenotypes we compared FUS protein level changes during MN differentiation between control and lines carrying mutations in FUS NLS. All of our tested lines showed that global FUS protein cellular levels were reduced upon MN differentiation but there was no significant difference in the amounts of this protein between *WT* and *FUS* mutant cells lines (Figure 1E, Suppl. Fig. 2A). Taken together, our previously characterized protocol of iPSC-based model of FUS-ALS showed normal differentiation into mature spinal MNs that acquire hallmark pathologies during their maturation and make a powerful model for pathophysiological studies of ALS mechanisms.

**Figure 1.**
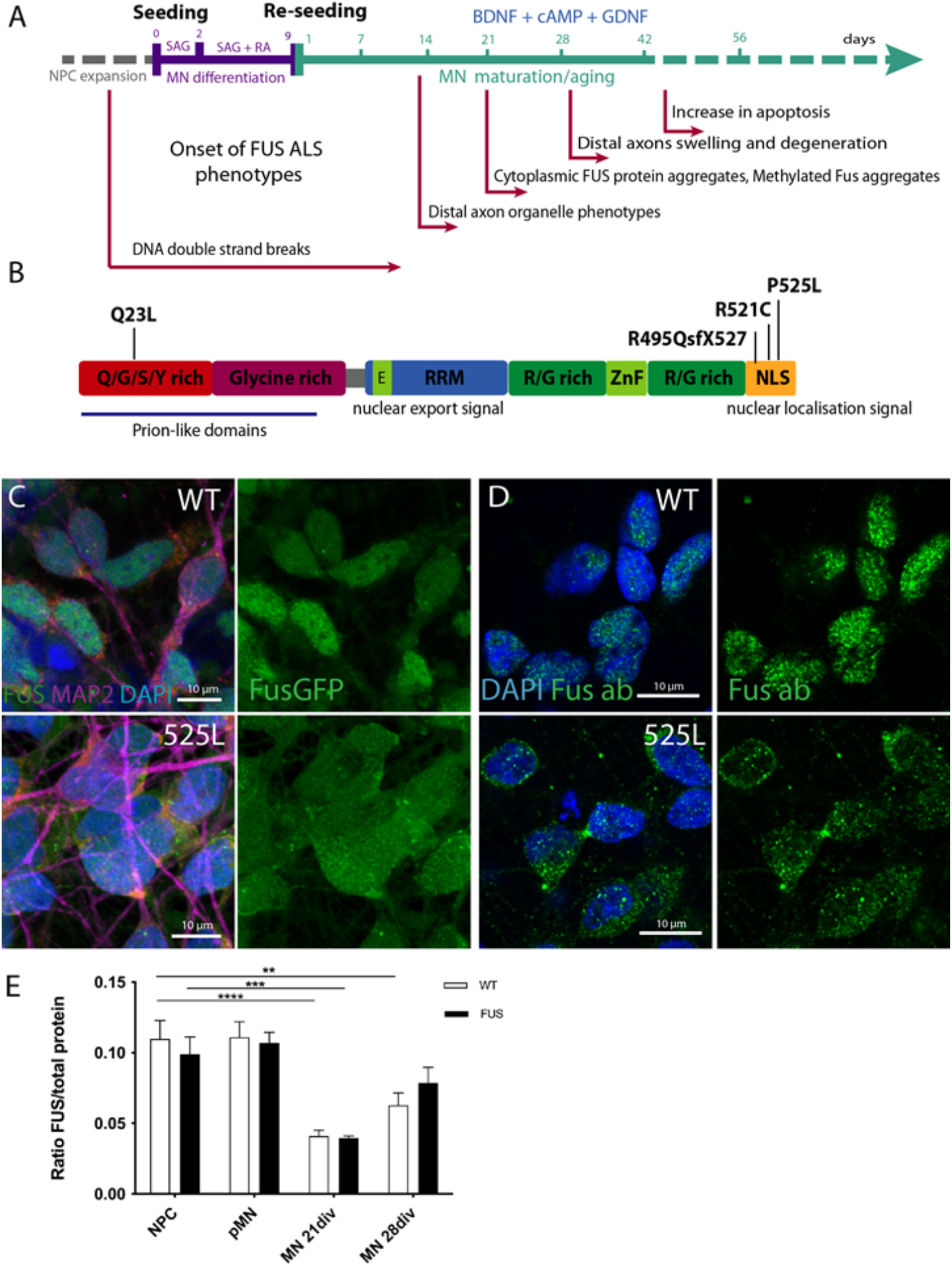
Generation of FUS ALS MNs from human iPSCs. **A.** Timeline and schematic outline of motoneuron differentiation protocol starting from neural precursor cells and undergoing sequential differentiation and maturation. Time points of the onset of previously reported FUS-ALS related phenotypes are indicated under the timeline. **B.** Schematic drawing of known protein domains of FUS and locations of respective FUS mutations analyzed in this study. **C.** Mature cultured MN showing mis-localization of mutant FUS^P525L^ protein from nucleus to the cytoplasm as seen by fluorescence imaging of FUS-GFP tagged isoforms or by immunostaining with FUS antibody. **D.** Bar chart quantification of western blot analysis demonstrating overall reduction of FUS protein levels during MN differentiation and maturation. Values were averaged from the individual measurements across all tested control and FUS mutant lines (see Suppl. Fig. 2A for individual lines evaluations). P values in this and all subsequent figure are displayed as * *p*<0.05, ** *p*<0.01, *** *p*<0.001. Scale bar 10 µm.

### 3.2. MN differentiation leads to an increase in a number of mitochondrial complex proteins and selected RNAs and is not affected by FUS mutations

Mature motoneurons, due to their morphology and function, are expected to have very high metabolic demands and high rates of oxidative phosphorylation (OXPHOS). It has been previously reported that MN differentiation is accompanied by a shift in their metabolic pathways from glycolysis to oxidative phosphorylation [10].We wanted to test if motoneuron differentiation is also accompanied by a general increase in metabolic rates/OXPHOS and revisit a potential role of *FUS*-ALS mutations during these changes. First, we tested if we could detect overall changes in mitochondrial material in our cell cultures during maturation. We collected cell protein extracts and mRNA samples at different stages of our motoneuron differentiation protocol: proliferating neural precursor cells (NPC), cells at the start of differentiation protocol (pre-motoneurons, pMN), as well as MN during maturation at days 7, 14, 21 and 28 after the start of the differentiation protocol (Fig. 2A). We then evaluated changes in the quantities of typical mitochondrial proteins and RNAs during differentiation. We compared three control and three independent lines carrying either P525L or R521C FUS ALS mutations (Table 1, Figure 2).

**Figure 2.**
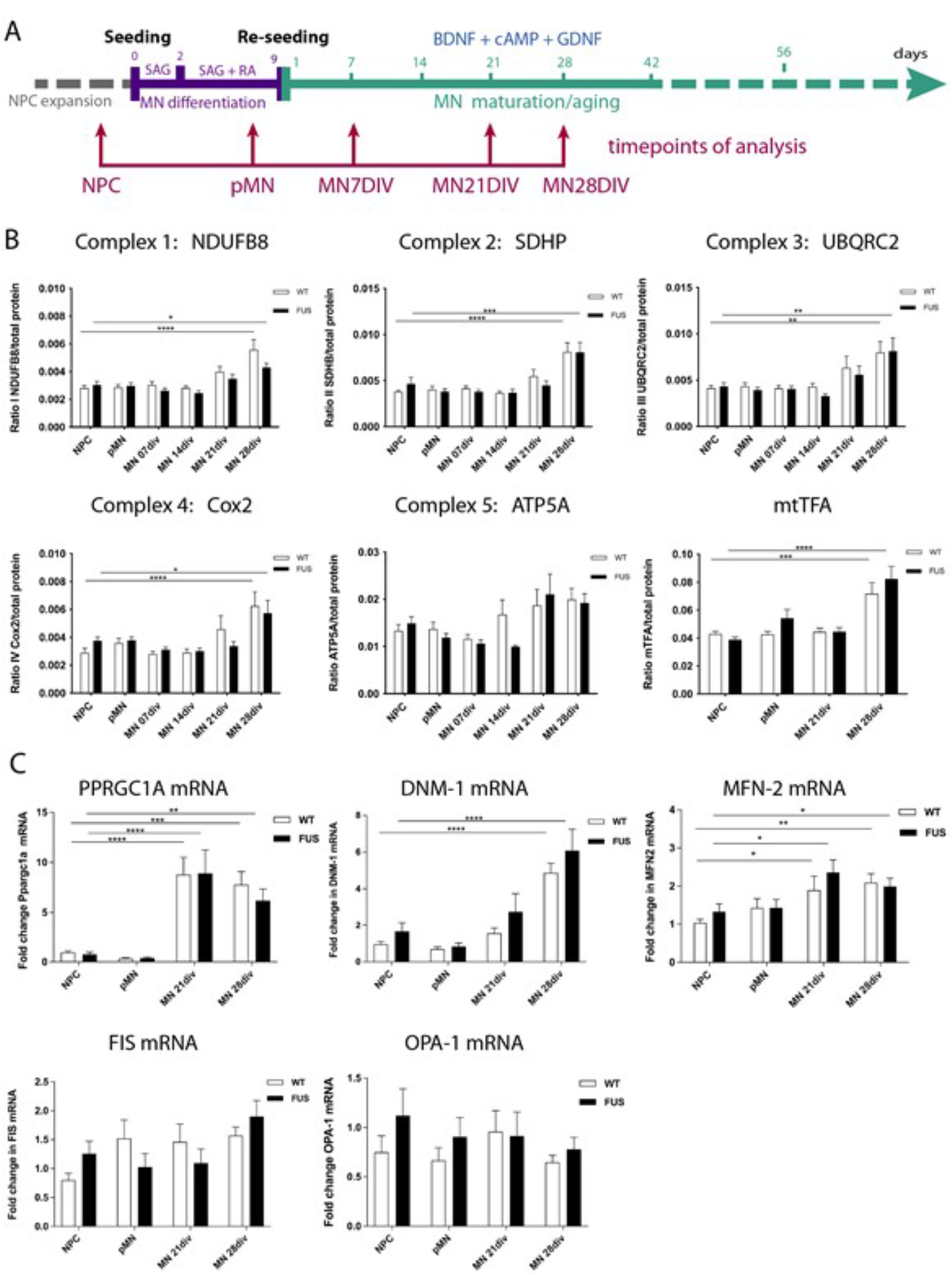
MN differentiation leads to an increase in mitochondrial complex proteins and mRNAs and is not affected by FUS. **A.**Timeline and schematic outline of motoneuron differentiation protocol showing timepoints at which cells were taken for analysis and are indicated by red arrows. **B.** Western blot analysis for selected components of the mitochondrial respiratory chain subunits I-V: NDUFB8, SDHB, UQCRC2, COX2, ATP5A, as well as mitochondrial transcription factor A(mtTFA) show increase in the relative amounts of these proteins (excluding ATP5A) in 28 day old MNs. **C**. qPCR evaluation of several mRNAs critical for maintenance and operation of the mitochondrial network like PPARGC-1α, Dnm1L and MFN-2 show an increase in the relative amounts by 21 or 28 days of MN maturation. Expression levels of OPA1 and FIS1do not change. **D.** Measurements of total amounts of mitochondrial DNA copy number (mtDNA) averaged across all tested WT and FUS lines. Presence of any FUS mutations did not significantly affect levels and amounts of any of the above tested proteins and mRNA (A-C). Values were averaged from the individual measurements across all tested control and FUS mutant lines (see Suppl. Fig 2B and Suppl. Fig 3). Significance was tested comparing cells at NPC stage to all other timepoints. * *p* < 0.05, ** *p* < 0.01, *** *p* < 0.001.

We assessed protein levels of single representative components from each of the mitochondrial respiratory chain subunits: 1) NADH dehydrogenase (ubiquinone) 1 beta subcomplex subunit 8 (NDUFB8), 2) succinate dehydrogenase (ubiquinone) flavoprotein subunit B (SDHB), 3) cytochrome b-c1 complex subunit 2 (UQCRC2), 4) cytochrome c oxidase subunit 2 (COX2), and 5) ATP synthase subunit alpha (ATP5A), as well as mitochondrial transcription factor A (mtTFA) (Fig 2). Four of the tested proteins showed a significant increase in MN by day 28 of maturation. ATP5A had a similar trend towards an increase but the p-values did not reach significance (Figure 2A).

In parallel to above mentioned increase in proteins of the mitochondrial complexes, we also detected a global increase in mRNA levels of Peroxisome proliferator-activated receptor gamma coactivator 1-alpha (PPARGC-1α), a transcriptional coactivator and the master regulator of mitochondrial biogenesis, Dnm1L, a mitochondria-associated, dynamin-related GTPase that mediates mitochondrial fission, and of mitofusin2 (MFN-2), a mitochondrial membrane protein that participates in mitochondrial fusion and contributes to the maintenance and operation of the mitochondrial network (Figure 2C). In contrast, there was no change during MN differentiation and maturation in the expression levels of the mitochondrial dynamin like GTPase OPA1 that regulates mitochondrial stability and energy output, as well as of FIS1, a component of a mitochondrial complex that promotes mitochondrial fission (Figure 2C). Of note, the presence of FUS mutations did not significantly affect levels and amounts of any of the above tested proteins and mRNA. Thus, MN differentiation and maturation leads to an increase in a number of mitochondrial proteins and RNAs, consistent with the increase in their expected metabolic rates and switch to oxidative phosphorylation, which was not affected by FUS-ALS causing mutations.

### 3.3. NAD^+^/NADH concentrations and redox ratio increases during MN differentiation and NAD^+^ concentrations increased in mature FUS-ALS MN

Concentrations of NAD^+^ and NADH, as well as NAD^+^ to NADH ratio are important parameters reflecting the general metabolic and redox state of different cell types during development, aging, and in a variety of human diseases, particularly in neurodegenerative disorders [70-72]. Increased NAD^+^/NADH ratio was reported to correlate with increased metabolic rates [73]. Furthermore, inhibition of mitochondrial electron transport chain activity decreases mitochondrial conversion of NADH to NAD^+^ and reduces the mitochondrial NAD^+^/NADH ratio [74]. Using cell extracts and a NAD^+^/NADH assay kit we analyzed how NAD^+^ and NADH concentrations and their ratio is changing upon differentiation and maturation of control and FUS ALS MN lines. Comparing averaged and individual values of all tested lines we observed an increase of NAD^+^/NADH concentrations and redox ratio upon differentiation and maturation of MN, with highest ratios detected in most mature MNs at day 28 (Figure 3A, Suppl. Fig. 4). Once again suggesting an overall increase in the expected metabolic rates and switch to oxidative phosphorylation in mature neuronal cells. As a positive control, we used treatment with FK866 to inhibit nicotinamide phosphoribosyltransferase, a key enzyme in the NAD^+^ and NADH biosynthesis from the natural precursor nicotinamide [75]. As expected, this treatment dramatically reduced NAD^+^, NADH levels as well as NAD^+^/NADH ratio in all MN.There was a significant increase in NAD^+^ concentration in mature FUS ALS MN that carry NLS FUS mutations. However, NADH concentration or NAD^+^/NADH redox ratio values were not significantly changed between control and FUS ALS MN (Figure 3A, Suppl. Fig. 4). This could mean that mitochondrial conversion of NADH to NAD^+^ is reduced in FUS ALS MN.

**Figure 3.**
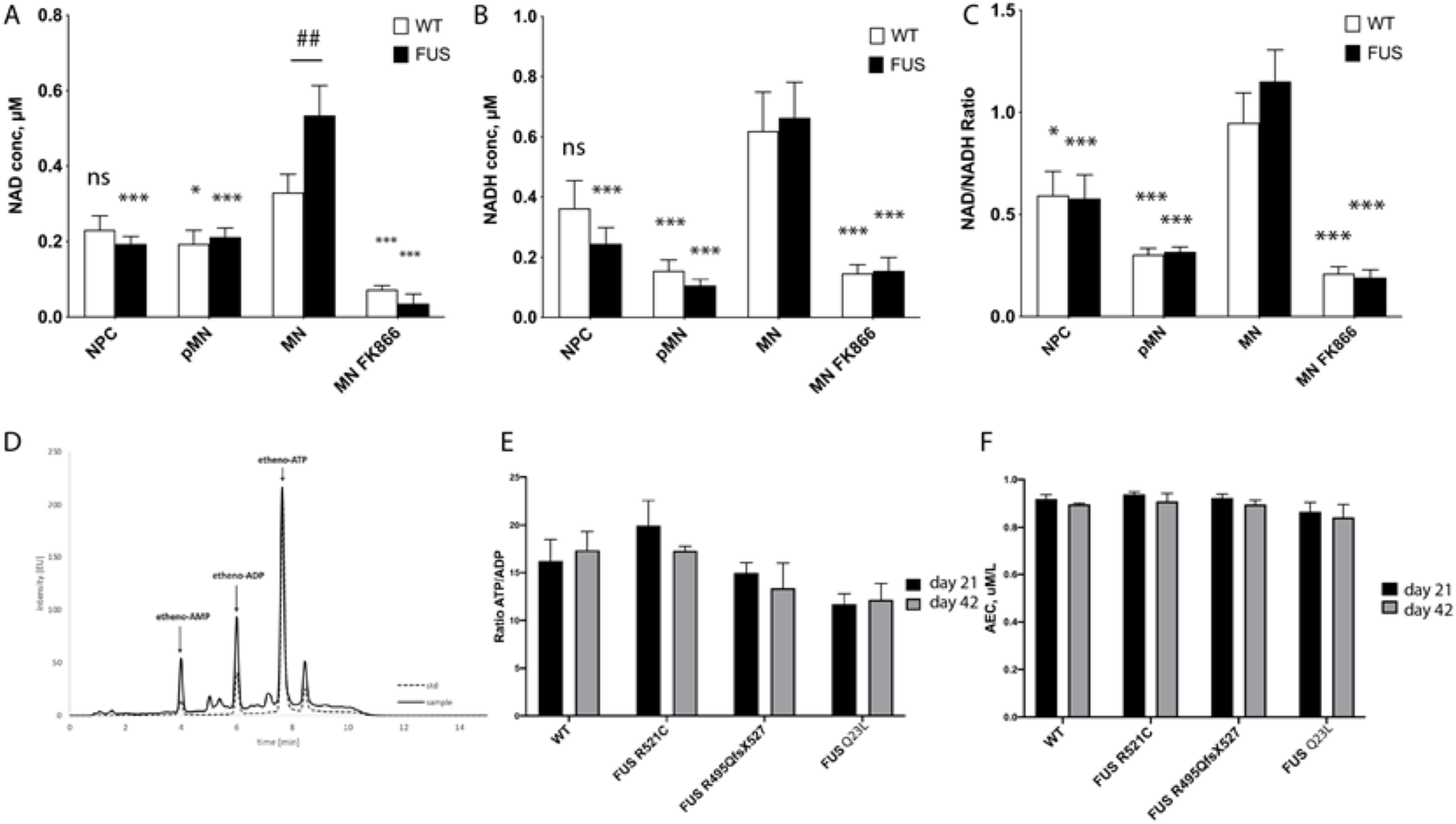
Evaluation of metabolic changes during MN differentiation of FUS-ALS cultured motoneurons. **A-C.** Measurements of NAD^+^ **(A)**, NADH concentrations **(B)** and their redox ratios **(C)** were averaged across all tested WT and FUS MN lines and displayed as bar graphs. Significance testing comparing values of mature MN to different time points of the same phenotype is shown with asterisks (*). Number signs (#) denotes significance testing between WT and FUS cells within the same time point. Significance testing two way Anova with Sidak method. For individual line measurements across all tested control and FUS mutant lines see Suppl. Fig 4. * *p* < 0.05, ** *p* < 0.01, *** *p* < 0.001, **D.** An example of typical chromatogram of adenine nucleotides in a whole cell motoneuron extract sample. **E-F.** Comparison of measured ATP/ADP concentration ratios **(E)** and adenylate energy charge (AEC) **(F)** in control and FUS-ALS mutants (R521C, R495QsfX527, Q23L) at two different maturation time points: day 21 and day 42 of MN maturation. We did not detect significant changes in in FUS ALS mutants at either time point.

### 3.4. Global measurements of adenosine nucleotides are not affected by FUS mutations

To further assess and compare metabolic rates and changes in FUS ALS MNs we decided to measure cytoplasmic levels of adenine nucleotides in the cellular lysates of mature MNs. Adenine nucleotides and adenosines form an important class of molecules that are central to both intra- and extracellular metabolic processes and are routinely used to evaluate metabolic states in the cell [54, 55, 76]. ATP/ADP ratio and adenylate energy charge (AEC) has been adopted to describe the energetic status of a cell or tissue preparation. We used 21 and 42 day old MN cell extracts in reversed-phase high-performance liquid chromatography (RP-HPLC) as a reliable, rapid simultaneous analytical determination of ATP, ADP and AMP concentrations (Figure 3D) [77]. We compared ATP/ADP ratio and adenylate energy charge (AEC) between *WT* and three different FUS ALS mutant lines (R521C, R495QsfX527 and Q23L). One striking observation, consistent with our NAD^+^/NADH measurements, was that we detected high ATP/ADP ratio in all of our MN cultures (up to 16-fold ratio). For reference, the ratio in hepatocytes, cells known to have high metabolic rates, was shown to be 6.6 [78]. These values yet again suggest high rates of OXPHOS occurring in mature MN. However, our measurements could not detect significant differences either in ATP/ADP cytoplasmic ratio or in AEC (Figure 3, C-D) of both day 21 and day 42 neuronal cultures carrying any of the tested FUS mutations. These data further support that FUS-ALS MNs have unchanged steady state adenine nucleotide concentrations which may indicate overall stable metabolic rates in our model ALS cell cultured MNs.

### 3.5. FRET-based ATP sensor allows metabolic measurements at a single cell level and compartmental resolution

The results of our measurements and similar published results [10] are surprising in the light of reports about mitochondrial defects, vesicle transport disruptions and hypoexcitability observed in cell culture, *in vivo* models and patient samples previously reported for FUS-ALS [13][14, 15][16, 25]. Several factors can potentially mask the expected phenotypes. Firstly, despite their robustness and efficiency, *in vitro* differentiation protocols produce highly enriched but still mixed-cell populations. Non-MN cells in the culture can potentially mask specific changes in affected cells, even though the switch to OXPHOS is typical for post mitotic neurons. Secondly, changes in diseased cells could be happening only in sub-compartments of neuronal cells, with axons potentially being most vulnerable. Furthermore, it is possible that to detect metabolic changes in unstressed cells one would require even longer cultivation times. We previously showed that neurodegeneration become apparent only after prolonged culture of cells [23, 65] and these observations go in line with etiology and slow development of ALS in patients.

To overcome above potential pitfalls and obtain live cellular level resolution of measurements we decided to use the FRET-based biosensor A-team to visualize ATP levels inside single living cells [59, 79]. A-team1.03 employs the ε subunit of the bacterial FoF1-ATP synthase as an ATP sensory domain sandwiched between two fluorescence proteins, CFP as the donor fluorophore present N-terminally and YFP as the acceptor at the C-terminus of the ε subunit [79]. In the presence of ATP the ε subunit binds to ATP and contorts, drawing the two fluorescence proteins closer to each other. Thus, ATP alters the fluorescent spectra of A-team by changing the FRET efficiency between CFP and YFP. The major limitation using intensity-based FRET measurement is an assumption that all observable donor molecules undergo FRET. This is usually not the case. This varying “unbound” fraction of donor molecules introduces considerable uncertainty to the measured FRET efficiency, making comparisons between experiments impossible [80]. To overcome this problem we used Fluorescence lifetime imaging (FLIM) [80-82]. The lifetime is determined by building up a histogram of detected fluorescence events. This reveals most commonly a multi-exponential fluorescence decay (Figure 4C). Numerical curve fitting renders the fluorescence lifetime and the amplitude (i.e., number of detected photons). Since FRET decreases the donor lifetime one can quantify the extent to which FRET occurs, provided the donor life-time without FRET is known. This donor lifetime *Tm* serves as an absolute reference against which the FRET sample is analyzed. Therefore, FLIM-FRET is internally calibrated – a property alleviating many of the shortcomings of intensity-based FRET measurements. As our reference donor lifetime we used lifetime of the CFP donor in a R122K/ R126K mutant isoform of ATeam1.03 that has no detectable ATP binding and therefore no ATP-dependent FRET events, but is otherwise identical to the normal sensor [59, 79]. For delivery of both ATeam1.03 and ATeam1.03 R122K/ R126K we prepared a lentivirus and infected differentiating neurons with the last final re-plating of pMN, which were then matured for further 21 day. Lower levels of virus load during infection allowed to infect small numbers of neurons, thus enabling easy identification of individual somato-dendritic and distal axonal compartments (Fig. 4A).

**Figure 4.**
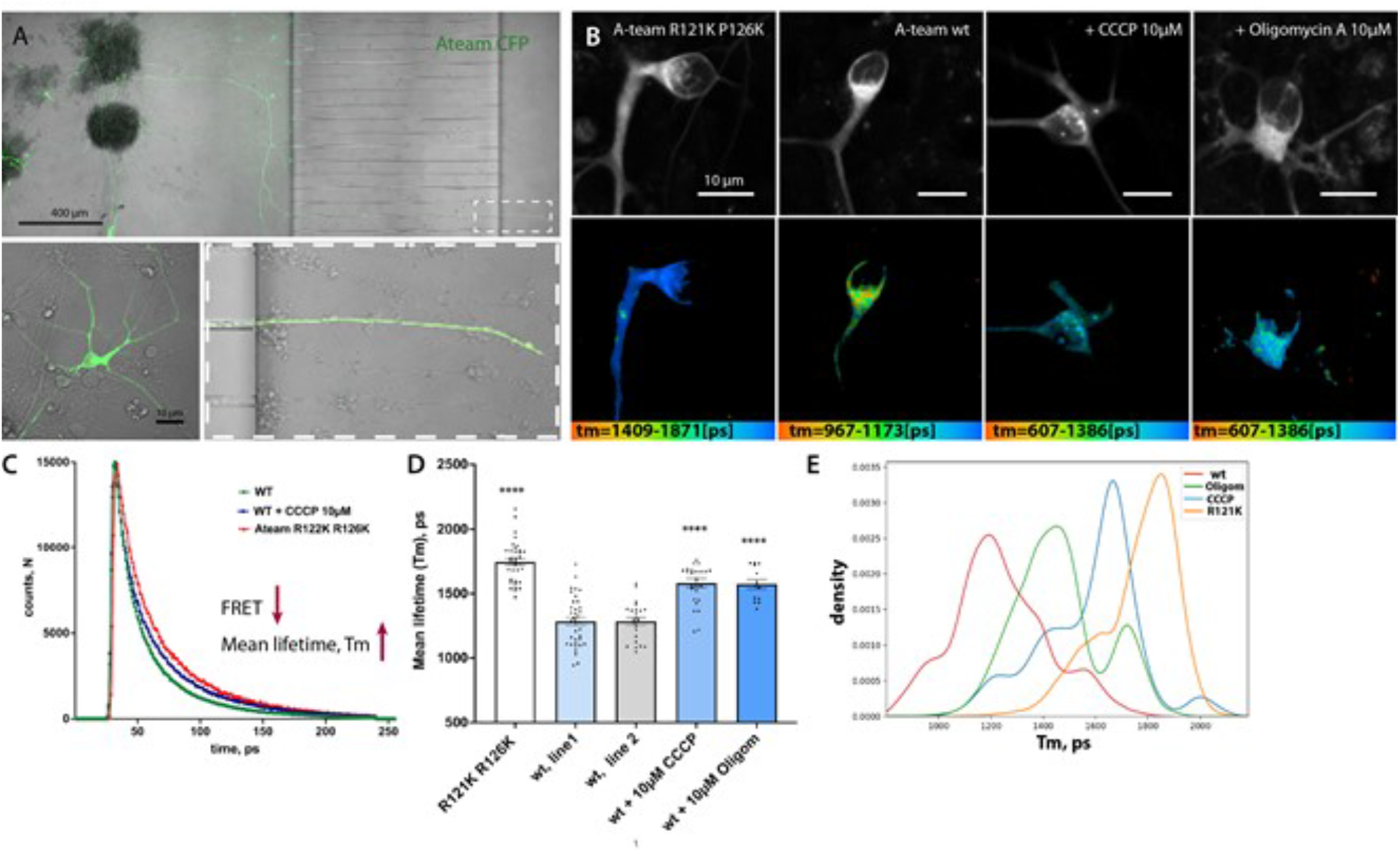
Lentiviral delivery of FRET-based ATP sensor A-team allows metabolic measurement at a single cell level and compartmental resolution. **A.** Low magnification of the microfluidic chamber demonstrating motoneurons transfected with lentivirus. Low to moderate transfection efficiency allows identifying single cell and measurements and tracing of individual soma, axonal and also distal axonal compartments within each chamber. **B.** Representative single images showing fluorescent intensity of donor (upper panel) and corresponding color-coded range of mean-lifetime FLIM measurements (lower panel) in transfected MN cells. **C.** Example single pixel FLIM decay model curves show shift towards increase of mean life time values in cells treated with mitochondrial blocker CCCP or transfected with a non-FRET control sensor A-team R121K R126K. **D.** Bar-graph chart showing changes in Mean lifetime measurements using A-team. Baseline FLIM measurements of the non-FRET donor fluorophore in a mutagenized A-team R121K R126K sensor, comparable average life-time measurements in two different untreated control cell lines and strong reduction of Tm values upon treatment with two different mitochondrial blockers CCCP (10 uM) and Oligomycin A (10 µM). Significance displayed is unequal variances T-test comparing wt1 to its corresponding treatments. **E.** Kernel density estimate (KDE) plot showing shifts in normalized distributions of Mean lifetime values per pixel in cells transfected with a non-FRET control sensor (R121K R126K) or with standard A-team without treatment (line wt1) or treated with two different mitochondrial blockers CCCP (10 µM) and Oligomycin A (10 µM).

As expected the mean lifetime (T_m_) of CFP-donor of the mutant non-FRET construct was significantly higher (i.e. low FRET levels) than that of the original functional A-team in two different *WT* control lines (*wt^1^* and *wt^2^*) (Tm(ATeam1.03 R122K/R126K) =1745+/-26.9, n=36 versus T_m_ *wt^.1^* =1283+/- 32.05, n=36 and T_m_ *wt^2^*=1284+/-30.05, n=24) (Fig. 4B, D, Suppl. Fig 5A). To make sure that these lifetime changes are indeed due to decreased ATP dependent change of FRET of the A-team sensor we applied two commonly used and well characterized inhibitors of final mitochondrial steps of OXPHOS, namely CCCP and Oligomycin A, to the MN. Both inhibitors caused a very strong increase in *Tm* in treated neurons: T_m_ ^CCCP^ =1581+/-34.43, n=26 and T_m_Oligom =1573 +/-36.71, n=13), an approximately 65% relative reduction in FRET efficiency in case of CCCP treatment (Fig. 4 B-D, Suppl. Fig. 5B-C). Thus, the A-team sensor performs well in living neurons and detects strong reduction of ATP levels upon mitochondrial inhibition of neuronal cells.

**Figure 5.**
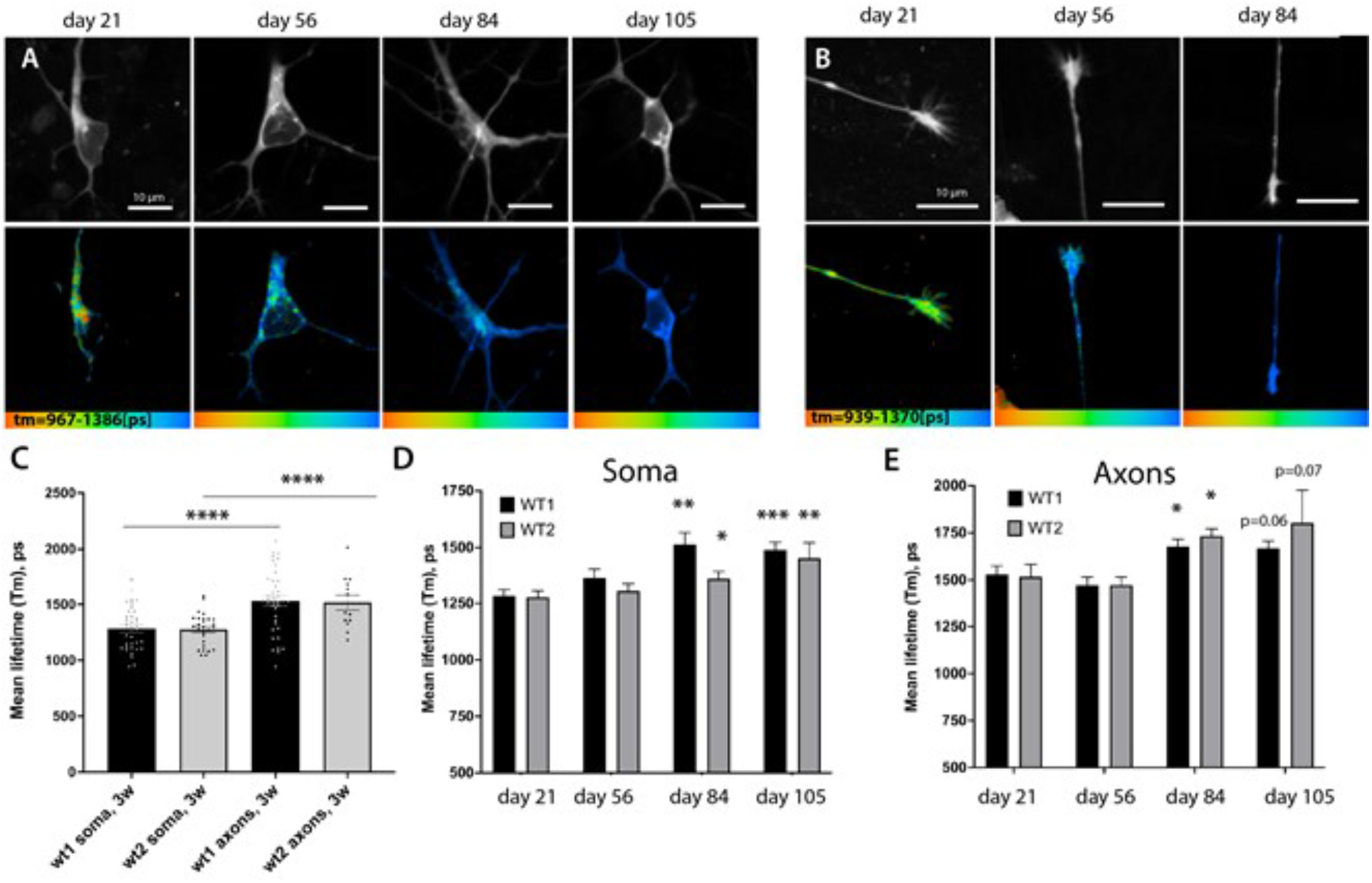
A-team FLIM revealed lower levels of ATP in distal axons and global slowing down of metabolism rates during prolonged culture of motoneurons. **A-B.** Representative images showing fluorescent intensity of donor (upper panel) and corresponding color-coded range of mean-lifetime FLIM measurements using A-team (lower panel). Images show different cell compartments, soma (A) and axons (B) in control MN cells at different time-points of cell culture: days 21, 56, 84 and 105 of maturation and aging. **C.** Bar graphs comparing average mean-lifetimes in soma and axons of two independent control lines of MN show strong significant reduction of relative ATP levels in distal axonal compartments at day 21. **D-E.** Prolonged culturing of MN leads to a significant global reduction of supposed ATP concentrations across neurons. Bar-chart showing increase in FLIM lifetimes of A-team donor fluorophore at various time-points post differentiation in soma (**D**) and distal axon (**E**) in two independent control lines upon long-term growth in culture. In **D** and **E** unequal variance T-test was used to compare the values between first timepoint 21 and later timepoints of the same line. * *p* < 0.05, ** *p* < 0.01, *** *p* < 0.001.

### 3.6. ATP levels in distal axons are significantly lower in comparison to soma

ALS is often reported as axonopathy since first symptomatic changes are associated with disruption of distal axons and neuromuscular junction (NMJ) muscle denervation [13, 83, 84]. Recent evidence also suggests high energy demands for NMJ and accumulation and anchoring of mitochondria in these sites [83]. We therefore wanted to measure the inherent differences between ATP levels in different sub-compartments of motoneurons to establish if distal axons are different in respect to soma. We compared average sensor lifetimes in soma and axons of two different patient-derived cell lines from healthy individuals (Figure 5 A-C). In both control lines FLIM lifetime values (T_m_) were significantly higher in the distal axons in comparison to soma: T_m_ ^soma^ *wt^1^* =1283+/-32.05, n=36 versus T_m_^axons^ *wt^.1^* =1530+/-44.99, n=39, p=0.0001; T_m_ ^soma^ *wt^2^*=1284+/-30.05, n=24 versus T_m_^axons^ *wt^2^*=1517+/-66.23, n=13, p=0.0003). These values suggest a strong relative reduction of ATP levels in distal axon and argue that distal axons have a significant disadvantage in energy supply. Analysis of Tm values distribution via frequency and sample size normalized kernel-density distributions (KDE) plot showed a clear shift of towards lower FRET values (Suppl. Fig. 5), therefore it is unlikely to be due to potentially smaller sampled area of axonal cytoplasm. Distribution curves also suggest a clear split between values with lower and higher T_m_, suggesting a less homogenous distribution of ATP in all conditions.

### 3.7. ATP levels across neuron drop during prolonged growth in culture

Slowing down of metabolism is a common hallmark in aging and is considered to be one of the critical risk factors that tip cellular homeostatic balance towards symptomatic onset of ALS and neurodegenerative diseases in general [70, 85]. For example, axoplasmic flows have been shown to significantly slow down with aging in rats and upon disruption of at least glycolytic metabolism [86, 87]. Although this slowing down of metabolism is a common assumption, precise measurements in specific cells have not been done extensively *in vivo* and especially in specific subsets of neuronal cells [85, 88].

We used MN cultures and *in vivo* sensor to monitor and detect whether any changes in metabolism could be detected in cells upon their prolonged culture. We found that using microfluidic chambers not only allows separation of compartments but also protects cultured MN from extra stress during media changes, increasing their survival time in comparison to larger cell formats. We have succeeded to robustly maintain MN cells for more than 105 days expressing the A-team sensor and continuously measured lifetime values both in soma and distal axons. Both our control lines showed a gradual increase of T_m_ values of A-team in the soma, and reaching a statistically significant reduction of relative ATP levels by day 84, where it plateaued for the rest of time-course (Figure 6A-C, Figure 7, Suppl. Fig. 7), with T_m_(wt1) (day 84)=1514+/-52.56, n=17, p=0.0003 and T_m(_wt2) (day 84)=1363+/-31.71, n=22, p=0.047). Axons, although starting from lower ATP levels baseline also showed a significant increase of sensor lifetimes by day 84 (Figure 5), suggesting that FRET efficiency and thus relative ATP levels are indeed reduced. Therefore, metabolic rates slow down across all regions of neuronal cells after a prolonged incubation in culture. Although, 105 days of growing of cells in culture unlikely corresponds to real aging, prolonged culturing of cells *in vitro* has been extremely useful to uncover some aspects of neurodegeneration mechanisms and it has been suggested that growing cells *in vitro* may put them under higher levels of stress and can mimic at least some aspects of ‘aging’ in cells [23].

**Figure 6.**
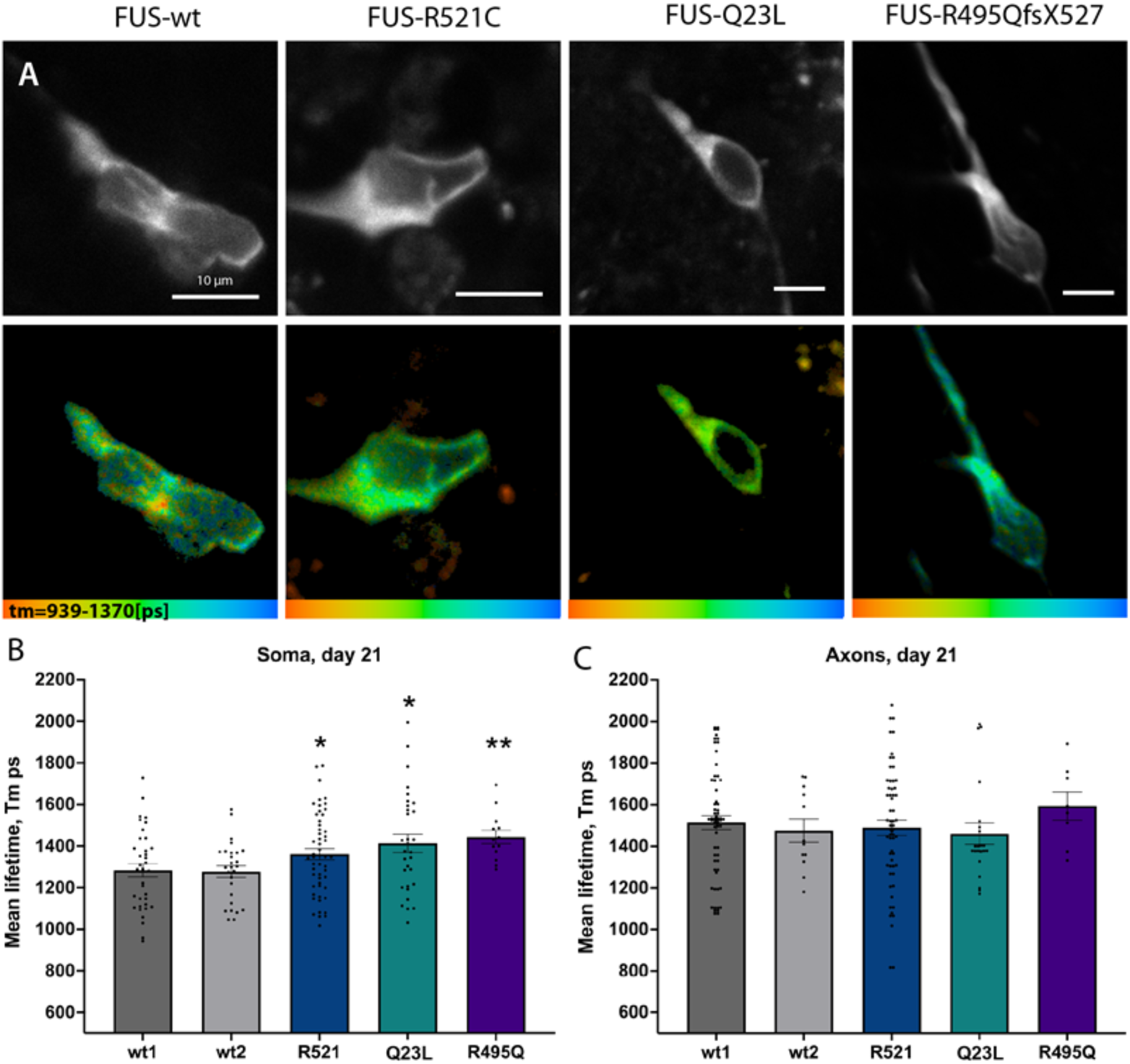
FUS ALS mutations led to early changes in ATP levels in cultured motoneurons. **A.** Representative single images showing fluorescent intensity of donor (upper panel) and corresponding color-coded range of the mean-lifetime from FLIM measurements (lower panel) in 21 day old mature control or FUS R521C, Q23L and R495QfsX527 mutant MNs. **B-C.** Bar graphs comparing average mean-lifetimes in soma **(B)** and axons **(C)**. Significances display differences between control line (wt1) and mutants and was assessed with unequal variance unpaired T-test. * *p* < 0.05, ** *p* < 0.01, *** *p* < 0.001.

**Figure 7.**
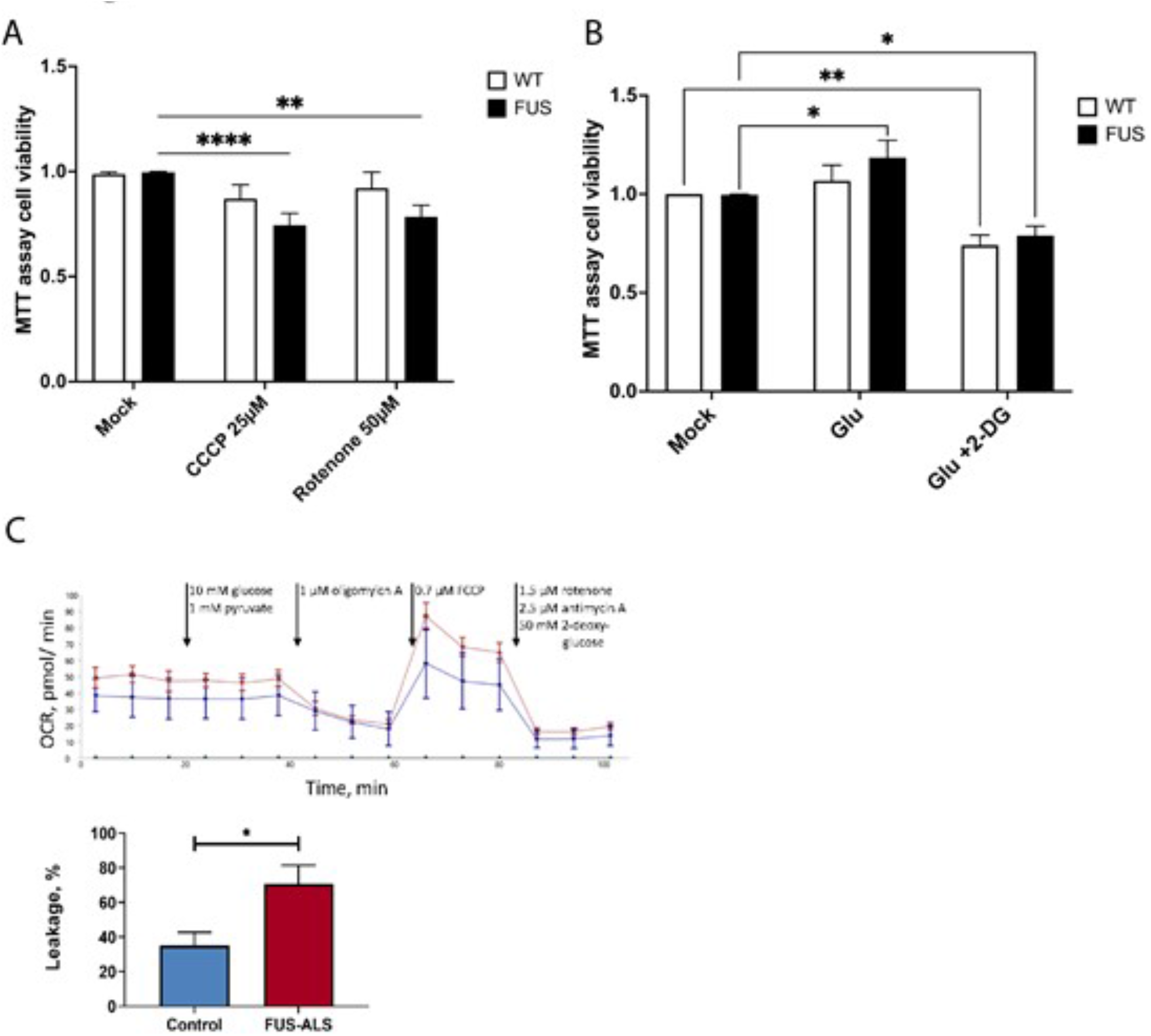
FUS ALS motoneurons have reduced viability upon additional mitochondrial stress and have an increased proton leakage. **A.** Bar graphs comparing cell survival of control and FUS ALS mutant MNs using MTT cell viability assay treated with mitochondrial inhibitors CCCP and Rotenone. Both mitochondrial inhibitors solely affected FUS mutant MN. * *p* < 0.05, ** *p* < 0.01, *** *p* < 0.001,Two-way ANOVA (Dunnet’s multiple comparison test). **B.** Glutamate stimulation of motoneurons and switching of metabolism to glycolytic pathways slightly improves baseline viability of FUS ALS MN, while blocking glycolysis by a utilization of 2-deoxy-d-glucose (2DG) reduces viability of both wt and FUS mutant MN. **C.** Example graph of seahorse measurement including the injections during the assay (arrows). **D.** FUS mutant neurons have an increased proton leakage through the inner mitochondria membrane. This result is calculated from OCR before and after oligomycin A injection. Bars represent mean+SEM; * represents p<0.01; n>7.

### 3.8. FUS ALS mutations lead to lower ATP levels in somas of cultured motoneurons

Having established a more sensitive and an *in vivo* way of measuring ATP levels we then decided to re-visit ATP levels measurements between control lines and three different FUSALS mutants available to us: two carrying mutations in nuclear-localization signal (NLS) (FUS^R521C^, FUS^R495QsfX527^) and one with a mutation in low complexity domain (LCD) (FUS^Q23L^). In support of previously suggested metabolism-related phenotypes we observed lower relative ATP concentrations (higher lifetime values of the sensor) in the somas of all FUS mutant motoneurons at day 21 of neuronal maturation (Fig. 6, Suppl. Fig 6C) with T_m_(FUS^R521C^)=1383+/-30.15, n=55, p= 0.0312, T_m_(FUS^Q23L^) =1413+/-43.27, n=31, p= 0.0169), T_m_(R495QsfX527) =1443+/-32.34, n=13, p=0.0073). Therefore our *in vivo* single-cell measurements detect early relative reduction of ATP in the soma of all tested FUS-ALS mutant motoneurons.

### 3.9. FUS MN have reduced viability upon further metabolic challenge

Lower levels of ATP at such early stage could facilitate accumulation of potential cellular defects associated with incorrect FUS function, affect its phase-transition mechanisms and overall make FUS ALS MNs less resilient to any additional external stress. To address the above hypotheses, we wanted to assess if FUS ALS MN are more vulnerable if challenged with additional metabolic stressors. We used MTT cell viability assay and treated three control and three FUS ALS MN cell lines with well characterized mitochondrial inhibitors: CCCP and Rotenone. Strikingly, two of the mitochondrial inhibitors at presented concentrations did not significantly affect viability of control cell lines but did clearly increase cell death in MN carrying FUS ALS mutations (Figure 7A, Suppl. Fig. 7).

We finally measured effect of glycolysis on cell survival. For this, we did the same tests together with either a block of glycolysis using 2-deoxy-d-glucose (2DG) or by inducing a shift of metabolism to glycolysis using glutamate [89-91]. Interestingly, using glutamate slightly improves baseline viability of FUS ALS MN. While blocking glycolysis reduces viability of control and mutant MN arguing that glycolysis is still essential for MN growth. Together these observations strongly support a role of FUS in maintaining metabolic states of MN via the regulation of mitochondrial oxidative phosphorylation and supporting their normal function, homeostasis and survival.

### 3.10. FUS mutations may cause proton leakage in mitochondrial membrane

For bulk metabolic analysis of patient specific MN we conducted seahorse measurement. Specific inhibition of different enzymes of glycolysis and respiratory chain was performed, while measuring the extracellular acidification rate (ECAR) as well as the oxygen consumption rate (OCR). An example graph is shown in Figure 7C. With the injection of port A (10 mM glucose, 1 mM pyruvate) the glycolytic increase over base line was measured but there was no difference observed between wt and FUS-mutant MN. Injection of a complex V inhibitor in port B (1 µM oligomycin A) allowed us to probe the cells for proton leakage from the intermembrane space to the matrix by measuring the OCR. There was a significant higher leakage observed in FUS mutant neurons when compared to *wt*. A decoupling agent was present in port C (0.7 µM FCCP). After its application the resulting OCR is indicative of the maximum respiratory capacity. However, there was no difference observed when wild type and disease neurons were compared. Lastly injection of port D (1.5 µM rotenone, 2.5 µM antimycin A and 50 mM 2-DG) was used to block glycolysis, complex I and complex II in order to measure absolute baseline of ECAR and OCR. ECAR and OCR measured during that period occur because of non-glycolytic acidification (e.g. Krebs cycle or breakdown of intracellular glycogen) and non-respiratory oxygen consumption (e.g. microsomal activities or cell surface oxygen consumption). We observed no differences between wt and FUS-mutant MN. Taken together seahorse showed no global phenotypes in oxidative phosphorylation and glycolysis but indicated a potential disturbance of the cristae structure by showing an increased proton leakage after oligomycin A application.

## 4. Discussion

To understand how to prevent, treat or even diagnose early such severe and fast-progressing neurodegenerative disease as ALS we need to understand and monitor changes that occur in vulnerable cells in the early disease course or even presymptomatic and in the context of development and aging. ALS and particularly FUS-ALS is caused by a disruption of the complex set of molecular mechanisms in motoneurons and their surrounding cells [2, 3][4]. In the majority of cases, however, changes occur incrementally and these mechanisms become prominent and critical only later in life [70, 88]. Aging itself is equally a complex and multifactorial process but is un-ambiguously accompanied by slowing down of metabolism and with it also disruption of protein-quality control systems [85]. Interestingly, these interlinked processes are especially critical for the etiology of many neurodegenerative diseases and not just FUS-ALS.

Abundant direct and indirect evidence points that metabolism does lie at the core of neurodegeneration and ALS [8-12][92, 93]. Indeed, work on at least FUS-ALS have reported defects in mitochondrial motility, morphology and function and would suggest a consequent reduction in the efficiency of oxidative-phosphorylation and metabolic stress for the highly-energy demanding motoneurons with particular vulnerability of their axons [6, 11][13][14, 15][16]. However, it is not trivial to monitor and detect specific metabolic changes in neurons over time. At least one recent report showed that metabolic changes do occur during MN differentiation but no differences in glycolytic or oxidative phosphorylation could be detected between control or FUS mutant motoneurons [10]. These observations are somewhat surprising given previously reported phenotypes, including mitochondria depolarization, reduced motility and fragmentation [23]. On the other hand, these results may reflect an inherent problem facing research of most neurodegenerative diseases, i.e. that initial critical changes at early stages of disease are subtle and incremental. Furthermore, these changes occur initially only in a specific neuronal sub-compartment. The reported mitochondrial defects manifest themselves first only in distal axons and are not initially visible in soma [13, 83, 84]. Keeping above considerations in mind we used previously established hiPSC-derived MN culture protocols and a range of techniques to re-visit measurements of metabolic rates in motoneurons. Our aims were to 1) increase sensitivity of our measurements; 2) asses changes in specific cell population (namely neurons) and compartments of neurons, such as soma and axons; 3) follow these measurements during differentiation, maturation and after prolonged culturing of the MN cells to detect early changes and monitor their progression.

Analysis of critical mitochondrial mRNAs, mitochondrial protein levels, high AEC ratio and NAD^+^/NADH concentrations and redox ratio all clearly showed a shift towards higher metabolic rates and shift towards OXPHOS in all MN upon differentiation and maturation and support expectations for the high energy demand of these highly specialized neuronal cells. FUS-ALS mutations did not affect most of these measurements and suggest that MN differentiation and growth occur normally in these cultures. We did however find first indication that FUS-ALS MN have changes in their metabolism and observed that global NAD^+^ concentrations were increased in all of the tested mutant lines. Although these changes did not significantly affect NAD^+^/NADH redox ratio, often used to estimate OXPHOS rates in cells, it showed that critical metabolite concentrations are affected. Maybe by a lack of mitochondrial conversion of NADH to NAD^+^ in FUS ALS MN. This motivated us to make use of more precise, live and single cell imaging tools to asses metabolism in single MN and their subcompartments.

Utilization of FRET sensors coupled with FLIM imaging to measure ATP levels showed a much more detailed and nuanced picture in both control and FUS-ALS MN. Firstly, single cell FLIM measurements highlight clear differences of ATP levels across cellular compartments in live cultured motoneurons. The somatic part of the cell has significantly higher ATP levels, which may reflect abundance of mitochondria and all accessory machinery in this compartment. Distal axons, on the other hand, from early on, have significantly lower concentrations of ATP. These observations can represent a reduction in mitochondrial number/density due to physical distance from the cell body and/or potential changes in the metabolic pathways utilized in axons. Recent data have shown mitochondria anchoring at NMJs and highlights their high energy demands [83]. Although our cultures did not contain NMJs, our measurements do emphasize that distance from the cell body or its compartmental metabolic specificity is a factor affecting energy availability in axons and puts them at a potentially higher risk of any further disruption of mitochondria and metabolism. The latter could explain why deterioration of NMJ and distal axons and muscle denervation is one of the first manifestations of ALS in humans and in multiple studies of ALS model systems [13, 83, 84]. Alternatively, distal axons could be presented with a different set of metabolites: either due to distance from the cell body soma, mode of metabolism in axons and/or differences in surrounding tissues. Importance of glycolysis to supplement energy supply in the synapse, as well as creatine metabolism and lactate supply from surrounding glia have been all reported to play an important role in axonal function and resistance to neurodegenerative processes [94-97]. All of these are however not part of our *in vitro* culture system presented here. Our data prompt further investigation of what distinguishes axonal compartment, which metabolites and pathways are involved and affected. We would like to further increase resolution of our measurements in both soma and axons and in the future monitor mitochondria in parallel to our FLIM measurements to get a further idea of ATP concentrations heterogeneity in relation and number of these organelles.

Most strikingly, single-cell FLIM measurements were able to detect a significant reduction of ATP levels in all three tested FUS-ALS mutant lines, including the most severe mutation of FUS R495QsfX527, known for a very early onset and aggressive subtype of ALS [66, 98]. Thus, metabolic changes in FUS-ALS do occur early in affected cells supporting previous observations and are likely to play an important detrimental role in ALS progression. Multiple mechanisms have been proposed for the FUS’ direct and indirect involvement of regulating metabolism, including aggregate accumulation of mitochondrial transcripts or direct binding to glycolytic enzymes and ATP synthase subunits [6, 11, 13-16]. Our data from cell viability assays and oxygen consumption rates measurement strongly argues that mutated FUS does affect function of mitochondria, increases proton leakage in the mitochondrial inner membrane and makes FUS-ALS MN much more sensitive to further inhibition of mitochondria and OXPHOS. This might also explain increased NAD^+^ levels by disturbing mitochondrial conversion of NADH to NAD^+.^. Whether these defects are due to reported FUS localization to mitochondria or direct binding to metabolic enzymes would need to be addressed in further studies. One further area of interest would be to characterize in much more detail changes in efficiency of glycolytic and OXPHOS pathways and their interaction in MN carrying ALS mutations. Increase in FUS-ALS MNs survival upon stimulation of glycolysis as well as clear shift in metabolic rates in all cell cultures after their prolonged cultivation suggests that some compensatory upregulation of glycolytic pathways could be beneficial to ALS MNs as was reported for in fALS models for *SOD1* and *TDP43* [99].

Interestingly, it was proposed that ATP concentration can regulate phase-transition properties of FUS and therefore potentially all transcription, translation and RNA metabolism processes regulated by such properties of FUS and similar proteins [38]. The fact that we detect lower energy levels in all FUS mutant model cell lines could be one of the factors responsible for gradual accumulating cell defects in disease cell. Cells can tolerate these changes initially but may become much more sensitive to any potential further stress after downregulation of metabolism that occurs in the cell and is proposed to happen in aging organism. Indeed, we and others observe much higher cell death rates, frequency of FUS aggregation and other ALS-related phenotypes in cells only after an extended time of cell culture and/or upon further externally added stress to the cell [65, 100] [23, 101]. Such properties of our FUS-ALS model cells would be in strong agreement with modern understanding progression of neurodegenerative diseases and their strong connection to underlying mechanism of aging and accumulating damage from external factors.

Our measurements cover only a very short timeframe and do not distinguish if this reduction was due to gradual change in metabolism in these cells or due to potentially high stress environment of the culturing neurons in 2D without their normal support network of cells. One would need to address both of these questions in further studies using metabolomic approaches or comparing OCR rates and ATP levels in *in vivo* model systems and compare these values in young and old organisms. However, to our knowledge there are very few examples of measurements in cell culture or *in vivo* models to demonstrate compartment-specific changes of metabolic rates with time. MN cultures could be therefore a powerful tool to study energy metabolism evolution with prolonged and ‘aged’ culture.

## 5. Conclusions

To be able to correctly diagnose and potentially develop treatments for ALS, we have to be able to monitor cellular changes in much more detail, with ideally live and sub-compartmental resolution under physiological conditions. Using FLIM-based non-invasive live cell imaging techniques that we have used, development of new sensors and expressing them in model organisms could become extremely beneficial for these purposes, particularly combined with the development of more advanced and physiological models to study ALS. Furthermore, our data confirms an important role of FUS in regulating mitochondrial and metabolic homeostasis in motoneurons and strongly supports the idea of developing potential new and proposed treatments that target energy metabolism, mitochondrial damage and oxidative stress.

## Supplementary Materials

Supplementary Figure 1. Initial characterization of the FUS-Q23L hiPSC line after fibroblast reprogramming; Supplementary Figure 2: Western blot analysis of FUS and mitochondrial proteins during FUS-ALS MN differentiation and maturation.; Supplementary Figure 3. Changes in expression levels of mRNAs important for mitochondrial function during FUS-ALS MN differentiation and maturation; Supplementary Figure 4. NAD^+^/NADH levels and redox ratio during MN differentiation of FUS-ALS cultured motoneurons; Supplementary Figure 5. Workflow of FLIM values extraction from single pixel and individual compartments ROI. Supplementary Figure 6: Comparison of individual pixel Tm values distributions in FLIM measurements between somatic and axonal compartments; Supplementary Figure 7: FUS ALS motoneurons have reduced viability upon additional mitochondrial stress.

## Author Contributions

Conceptualization, V.Z. and A.H.; methodology, V.Z., A.P. H.G., M.M, J.J., A.D., B.S and A.H.; validation, V.Z., A.P; J.J. formal analysis, V.Z, P.M.A. H.G..; investigation, V.Z. A.P, M.M, J.J. A.D., H.G,; resources, A.H., T.B, J.S, A.S., S.R, E.Z., C.D, J.S, P.M.A; data curation, V.Z., A.P, H.G., M.M; S.R. writing—original draft preparation, V.Z., H.G., S.R., J.J. and A.H.; writing—review and editing, all authors; visualization, V.Z., A.P, H.G.; supervision, A.H.,A.S., S.R, C.D., E.Z.; project administration, A.H.; funding acquisition, A.H., S.R., C.D, E.Z. All authors have read and agreed to the published version of the manuscript.

## Institutional Review Board Statement

The performed procedures were in accordance with the Declaration of Helsinki (WMA, 1964) and approved by the Ethical Committee of the Technische Universität Dresden, Germany (EK 393122012 and EK 45022009) and Nr 94-135 (to perform genetic research including for FUS) Nr 2018-494-32M (to perform in vitro lab studies on cell lines derived from patients with ALS fo Umea University.

## Informed Consent Statement

Written informed consent was obtained from all participants including for publication of any research results.

## Data Availability Statement

All data is presented in the manuscript

## Acknowledgments

We deeply thank the patients and probands taking part in this study. We acknowledge the great cell culture help of Anett Böhme and Katja Zoschke. We thank Tony Hyman lab for sharing plasmids for the A-team sensor constructs. The Light Microscopy Facility (LMF) of CMCB (Center for Molecular and Cellular Bioengineering, Technische Universität Dresden) provided excellent support for all live imaging experiments. We acknowledge Horst Wallrabe, Karsten Siller and the Keck Center for cellular imaging at UVA for the help with FLIM data analysis and validation and use of their FLIM analyzer tools and FIJI plugins (PI: AP; NIH OD016446). Vivek Khatri and Riya Verma for help with manual tracing of ROI during FLIM data analysis.

---

**Supplementary Figure 1.**
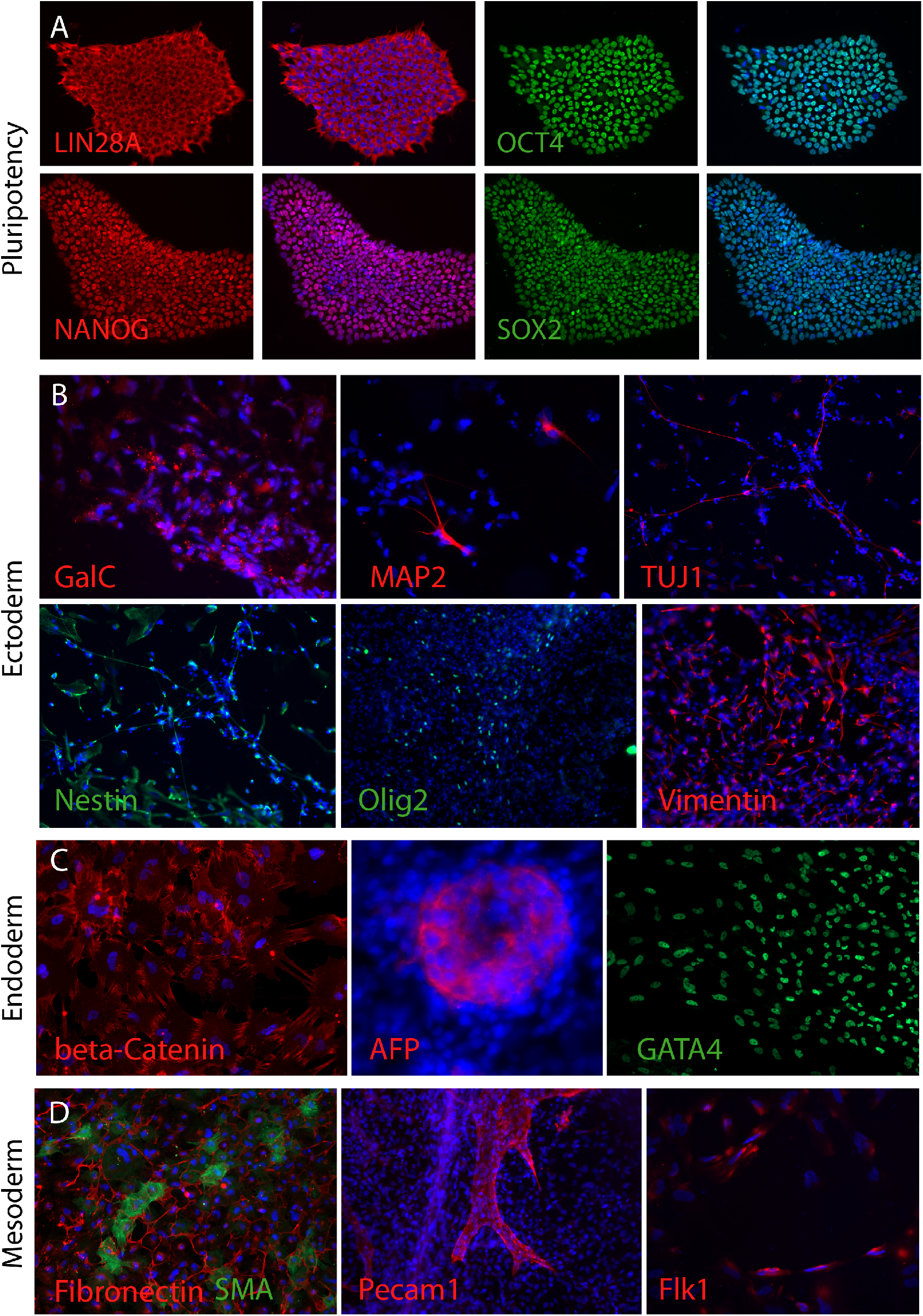
Initial characterization of the FUS-Q23L hiPSC line after fibroblast reprogramming. **A.** hiPCs derived from patient fibroblasts were tested for presence of pluripotency markers by immunostainings for Nanog, Oct4, Sox2 and Lin24A **B-D.** FUS-Q23L hiPSC colonies were able to differentiate into all three germ layers as shown by immunostainings for typical markers for Ectoderm **(B),** Endoderm **(C)** and Mesoderm **(D).**

**Supplementary Figure 2.**
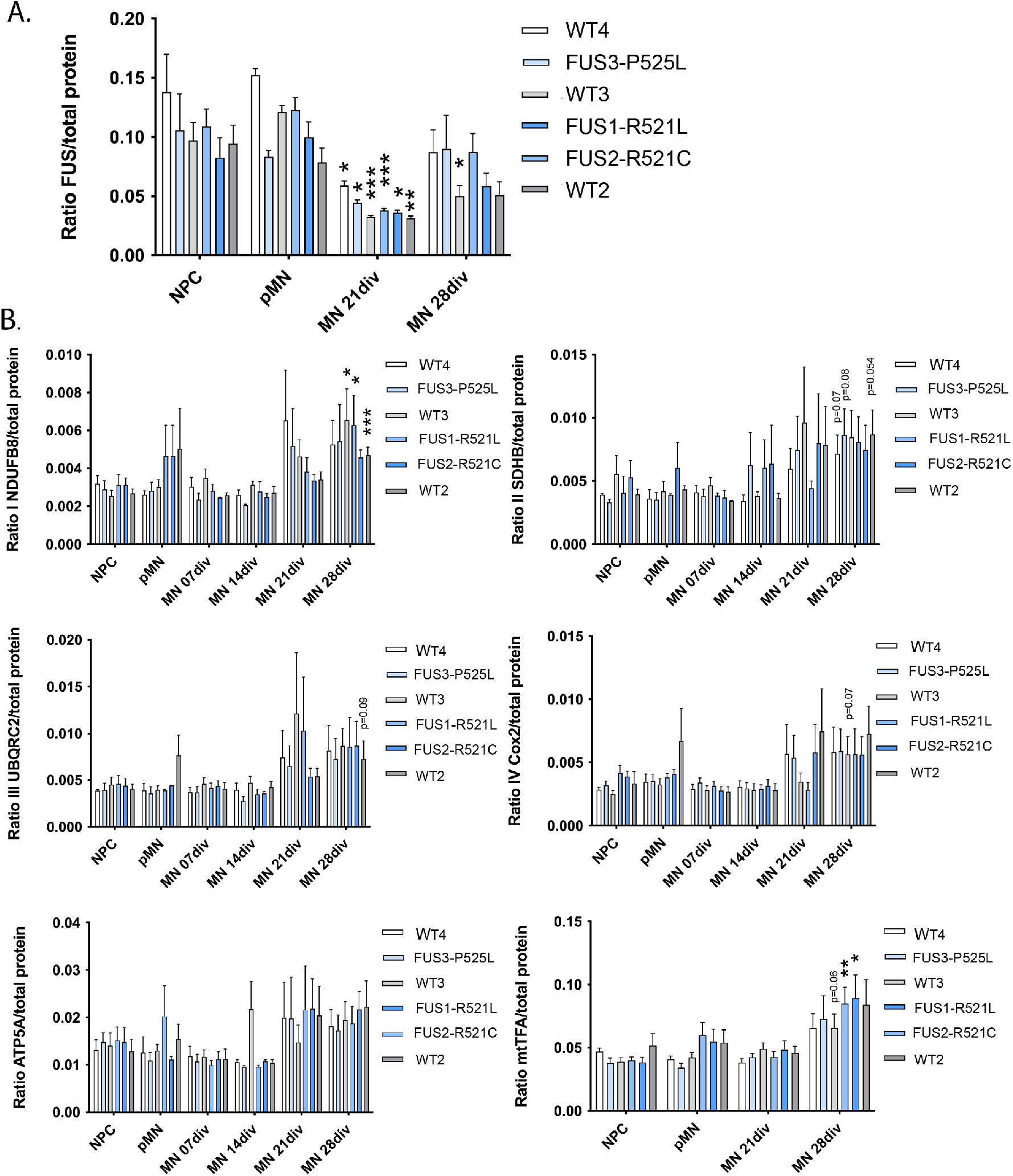
Western blot analysis of FUS and mitochondrial proteins during FUS-ALS MN differentiation and maturation. **A.** Quantification of western blots for FUS protein levels during MN differentiation and maturation in FUS-ALS lines. **B.** Quantification of western blots for selected components of the mitochondrial respiratory chain subunits I-V: NDUFB8, SDHB, UQCRC2, COX2, ATP5A, as well as mitochondrial transcription factor A(mtTFA) show increase in the relative amounts of these proteins (excluding ATP5A) in 28 day old MNs in FUS-ALS lines. Asterisks show significance comparing values between different stages of differentiation within the same line.

**Supplementary Figure 3.**
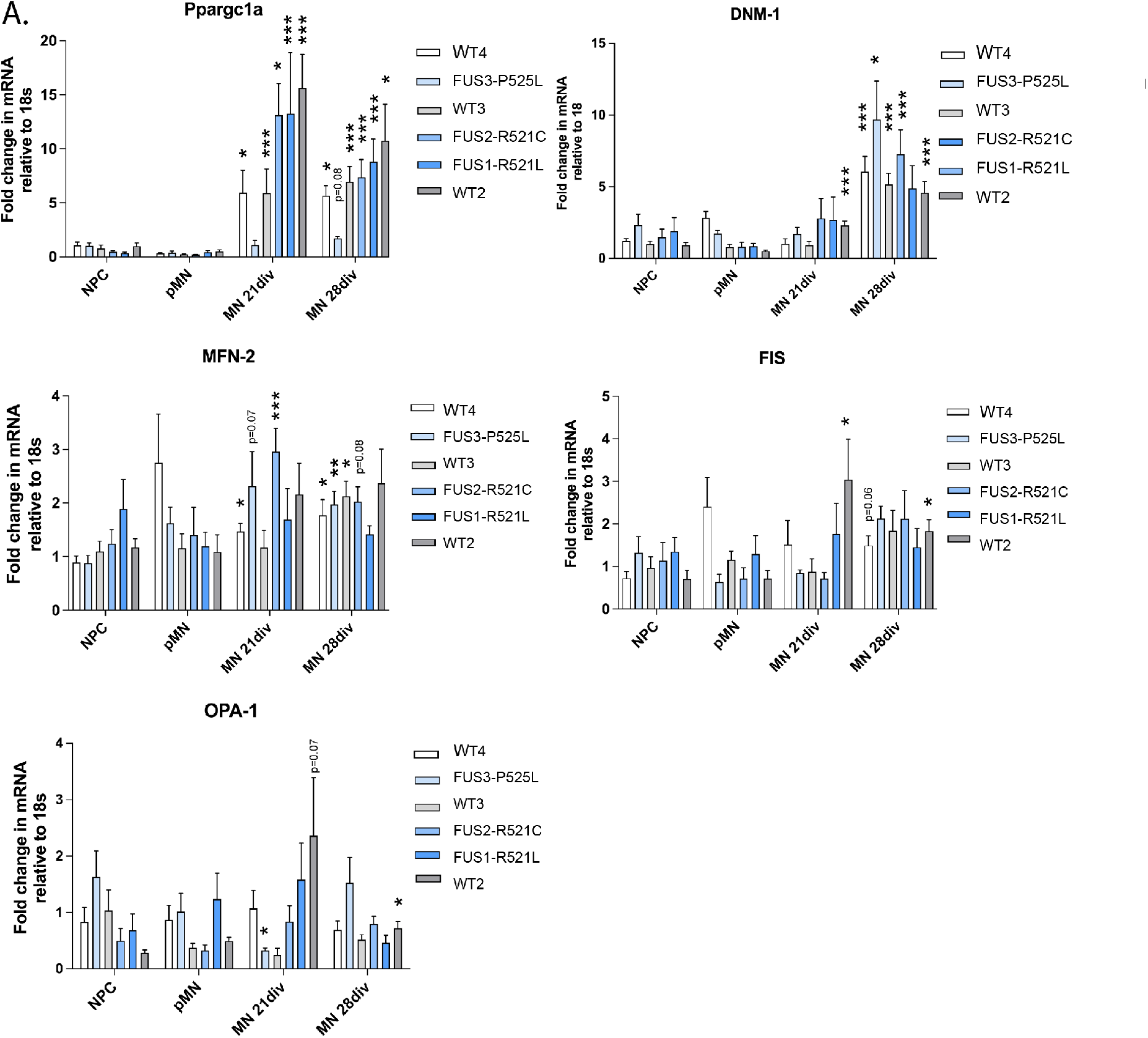
Changes in expression levels of mRNAs important for mitochondrial function during FUS-ALS MN differentiation and maturation. **A.** qPCR quantification of relative expression levels in several mRNAs critical for maintenance and operation of the mitochondrial network: PPARGC-1α, Dnm1L, MFN-2, OPA1 and FIS1 during MN differentiation and maturation in FUS-ALS lines. Asterisks show significance comparing values between different stages of differentiation within the same line.

**Supplementary Figure 4.**
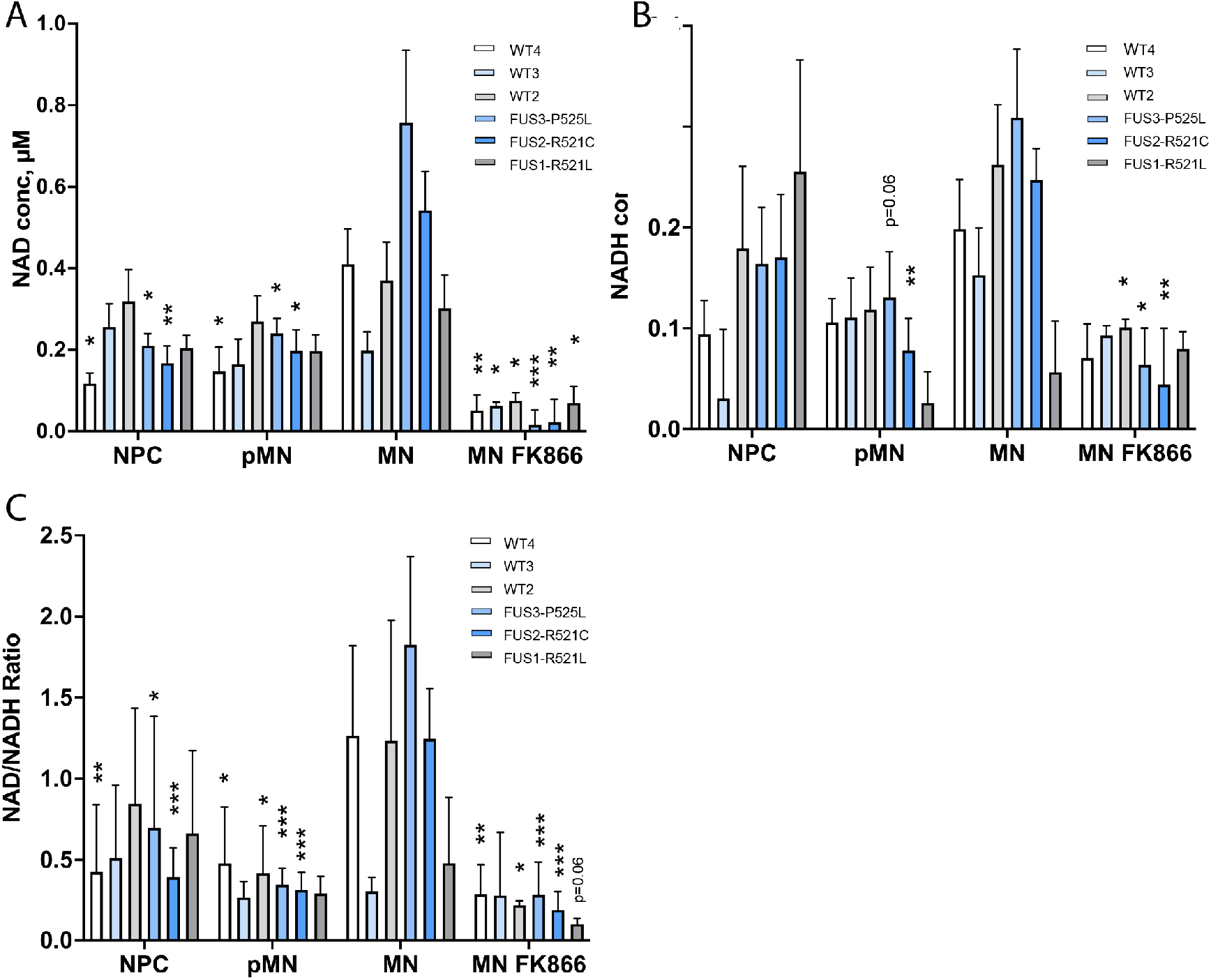
NAD+/NADH levels and redox ratio during MN differentiation of FUS-ALS cultured motoneurons. **A-C** Measurements of NAD^+^ **(A)**, NADH **(B)** concentrations and their redox ratios **(C)** across all tested individual control and FUS mutant lines at different stages of MN differentiation. Significances are shown as * between MN stage vs other stages or treatment within the same MN

**Supplementary Figure 5.**
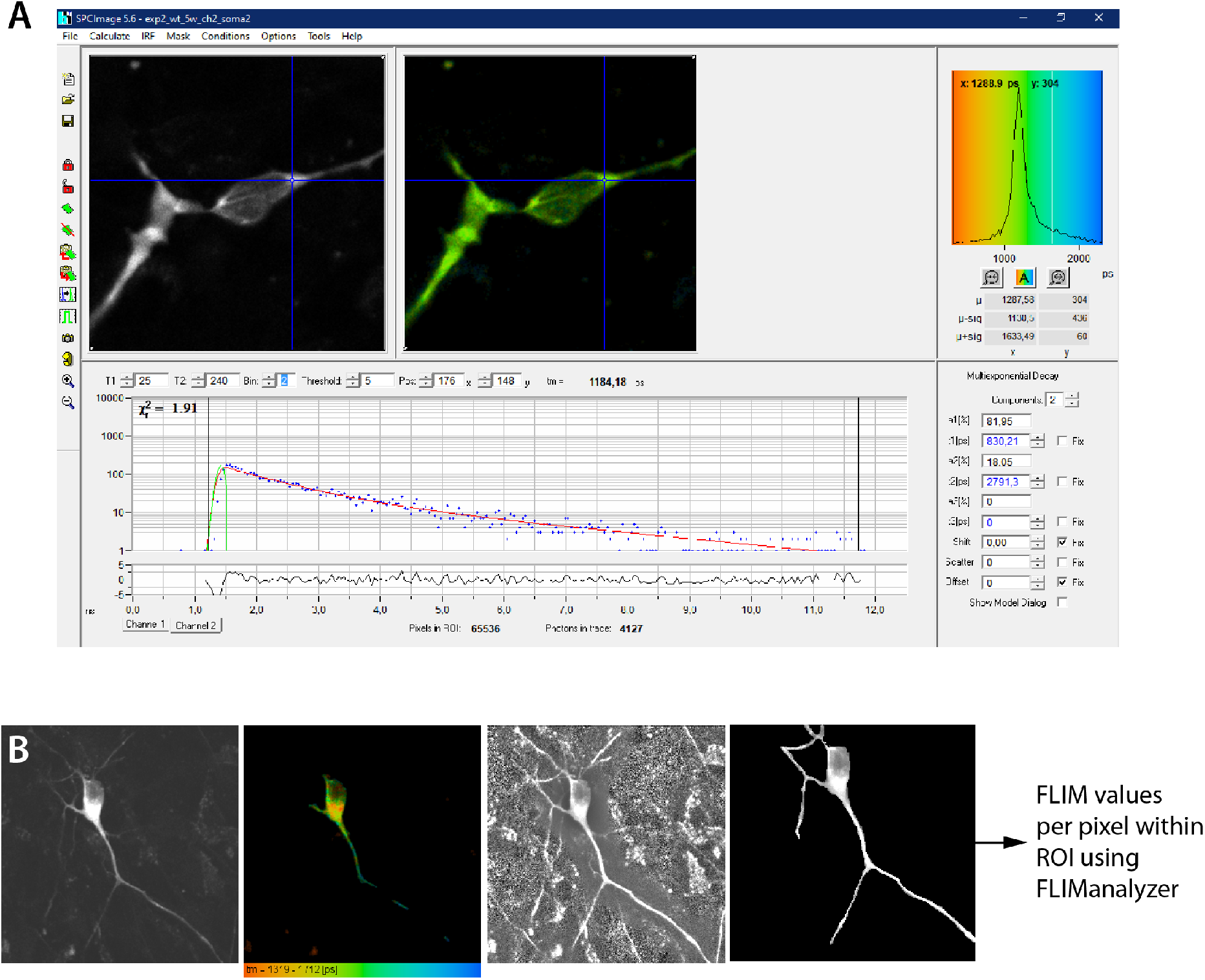
Workflow of FLIM values extraction from single pixel and individual compartments ROI. **A.** Snapshot of SPCImage software window with an example curve fitting for single pixel FLIM measurements. Modelling is based on two components incomplete model fitting. **B.** Fluorescence donor (CFP) intensity signal, color coded FLIM values per pixel, and contrast-guided selection of ROI for data extraction to exclude nuclei and background noise measurements.

**Supplementary Figure 6.**
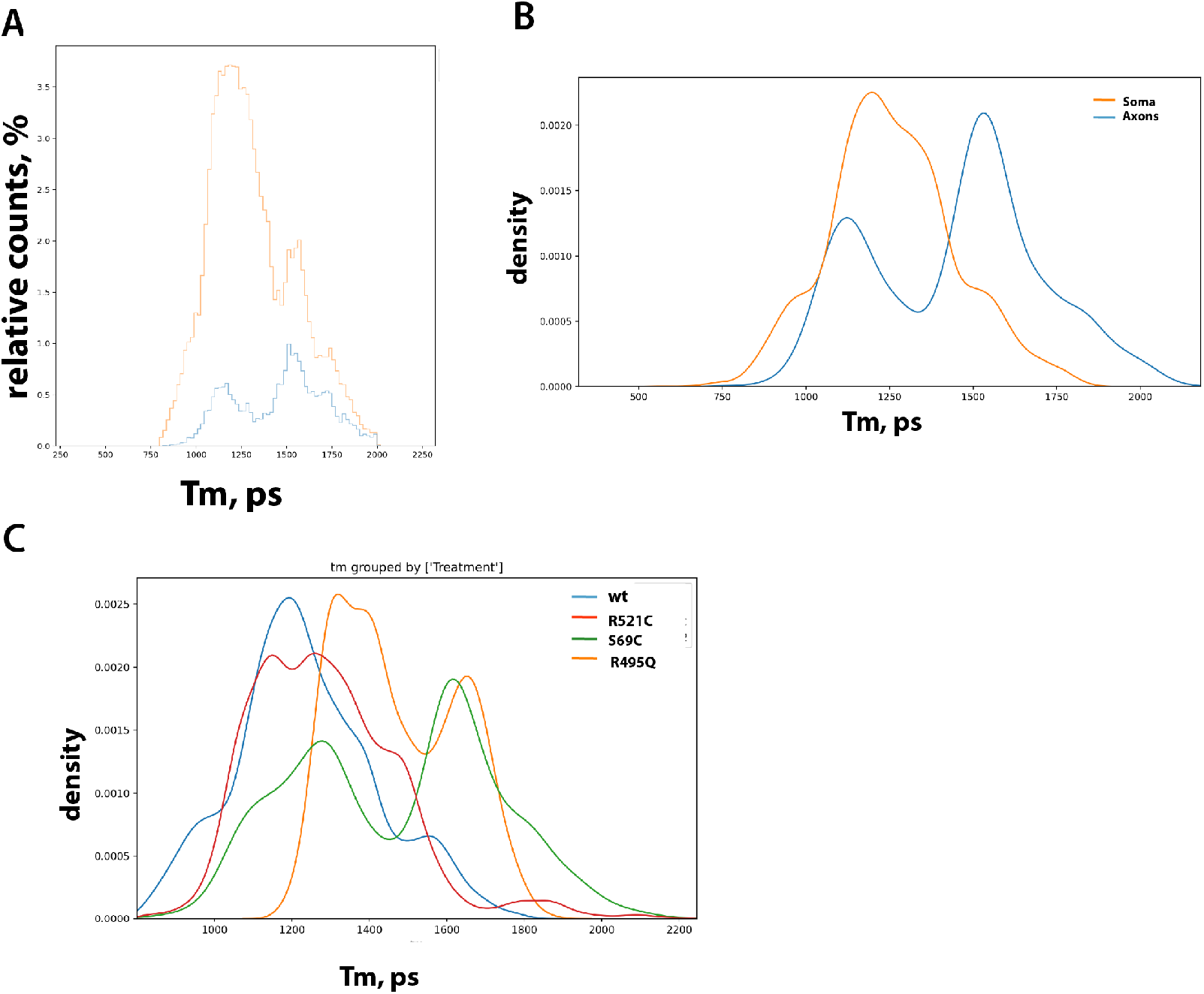
Comparison of individual pixel Tm values distributions in FLIM measurements between somatic and axonal compartments. **A.** Frequency distribution histogram **(A)** and normalized KDE distribution plot **(B-C)** comparing Tm values of somatic and axonal compartments **(B)**, as well as comparing control (wt1) and mutant FUS MN.

**Supplementary Figure 7.**
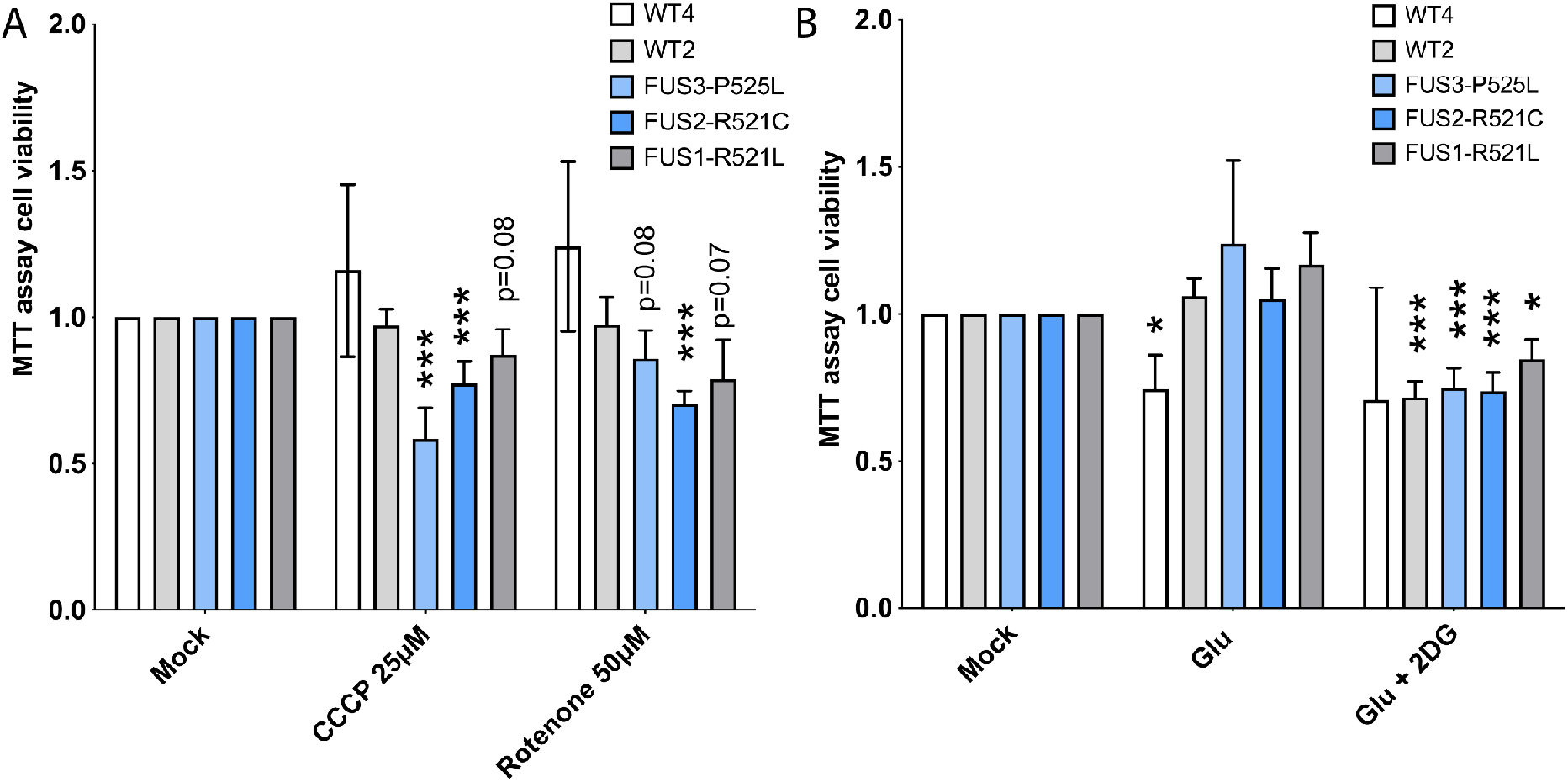
FUS ALS motoneurons have reduced viability upon additional mitochondrial stress. **A.** Bar graphs comparing cell survival of individual control and FUS ALS mutant MNs using MTT cell viability assay treated with mitochondrial inhibitors CCCP and Rotenone. **B.** Bar graphs comparing cell survival of individual control and FUS ALS mutant MNs using MTT cell viability assay after glutamate stimulation or blocking glycolysis with 2-deoxy-d-glucose (2DG).

