## Supplementary figures for "Live cell imaging of ATP levels reveals metabolic compartmentalization within motoneurons and early metabolic changes in *FUS* ALS motoneurons"

Supplementary Figure 1.

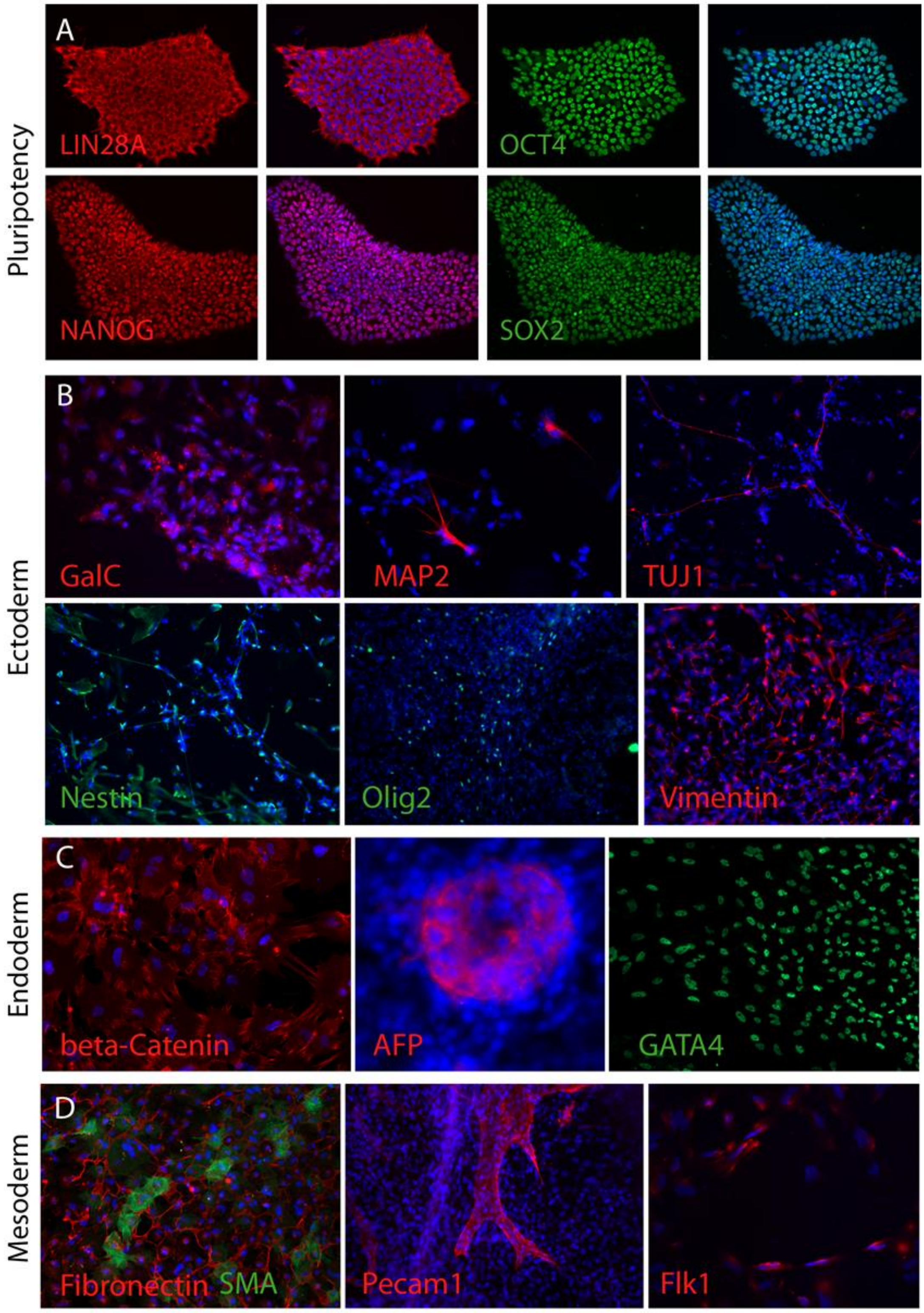


**Supplementary Figure 1. Initial characterization of the FUS-Q23L hiPSC line after fibroblast reprogramming. A.** hiPCs derived from patient fibroblasts were tested for presence of pluripotency markers by immunostainings for Nanog, Oct4, Sox2 and Lin24A **B-D.** FUS-Q23L hiPSC colonies were able to differentiate into all three germ layers as shown by immunostainings for typical markers for Ectoderm **(B),** Endoderm **(C)** and Mesoderm **(D).**

Supplementary Figure 2.

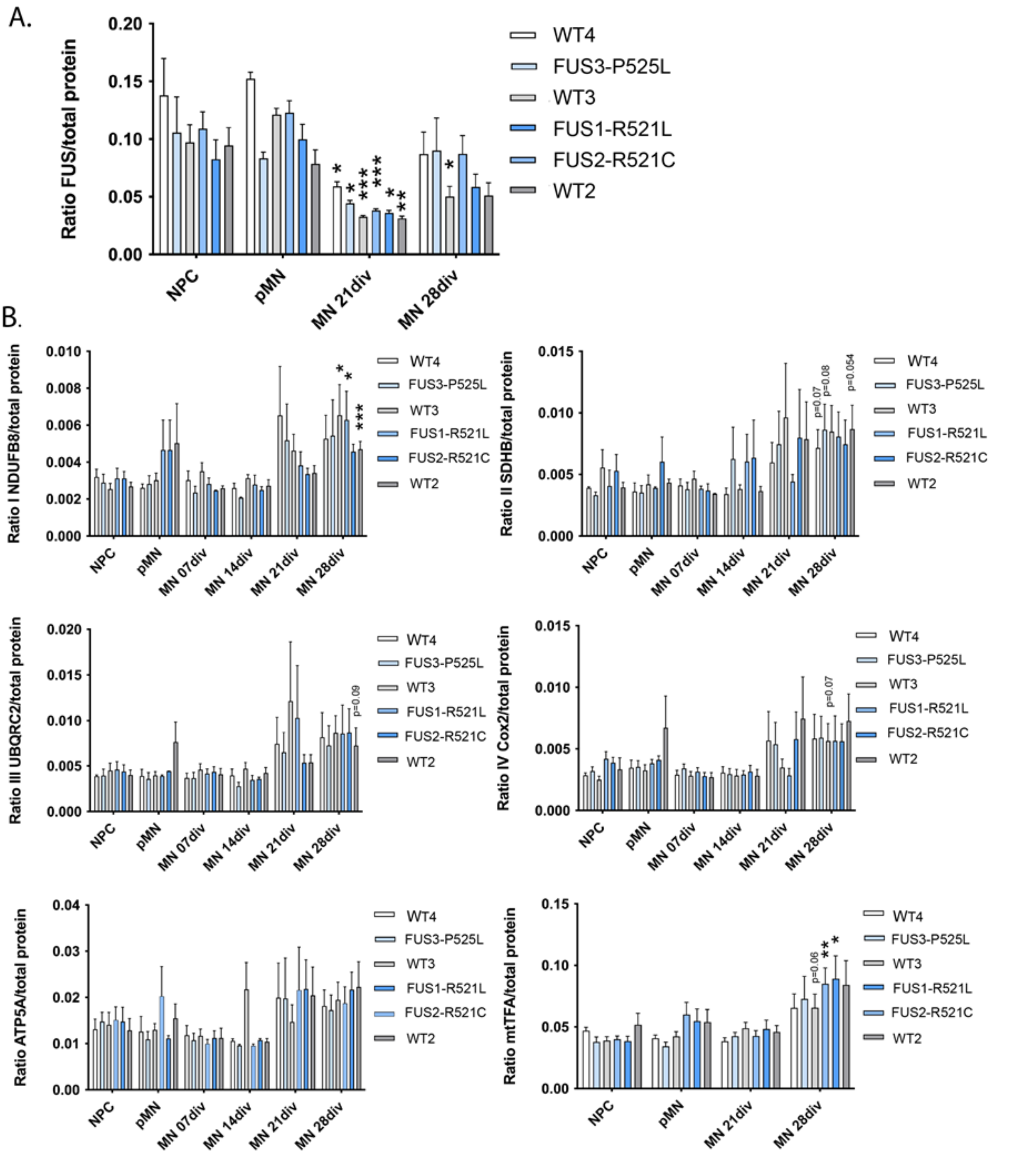


**Supplementary Figure 2. Western blot analysis of FUS and mitochondrial proteins during FUS-ALS MN differentiation and maturation. A.** Quantification of western blots for FUS protein levels during MN differentiation and maturation in FUS-ALS lines. **B.** Quantification of western blots for selected components of the mitochondrial respiratory chain subunits I-V: NDUFB8, SDHB, UQCRC2, COX2, ATP5A, as well as mitochondrial transcription factor A(mtTFA) show increase in the relative amounts of these proteins (excluding ATP5A) in 28 day old MNs in FUS-ALS lines. Asterisks show significance comparing values between different stages of differentiation within the same line.

Supplementary Figure 3.

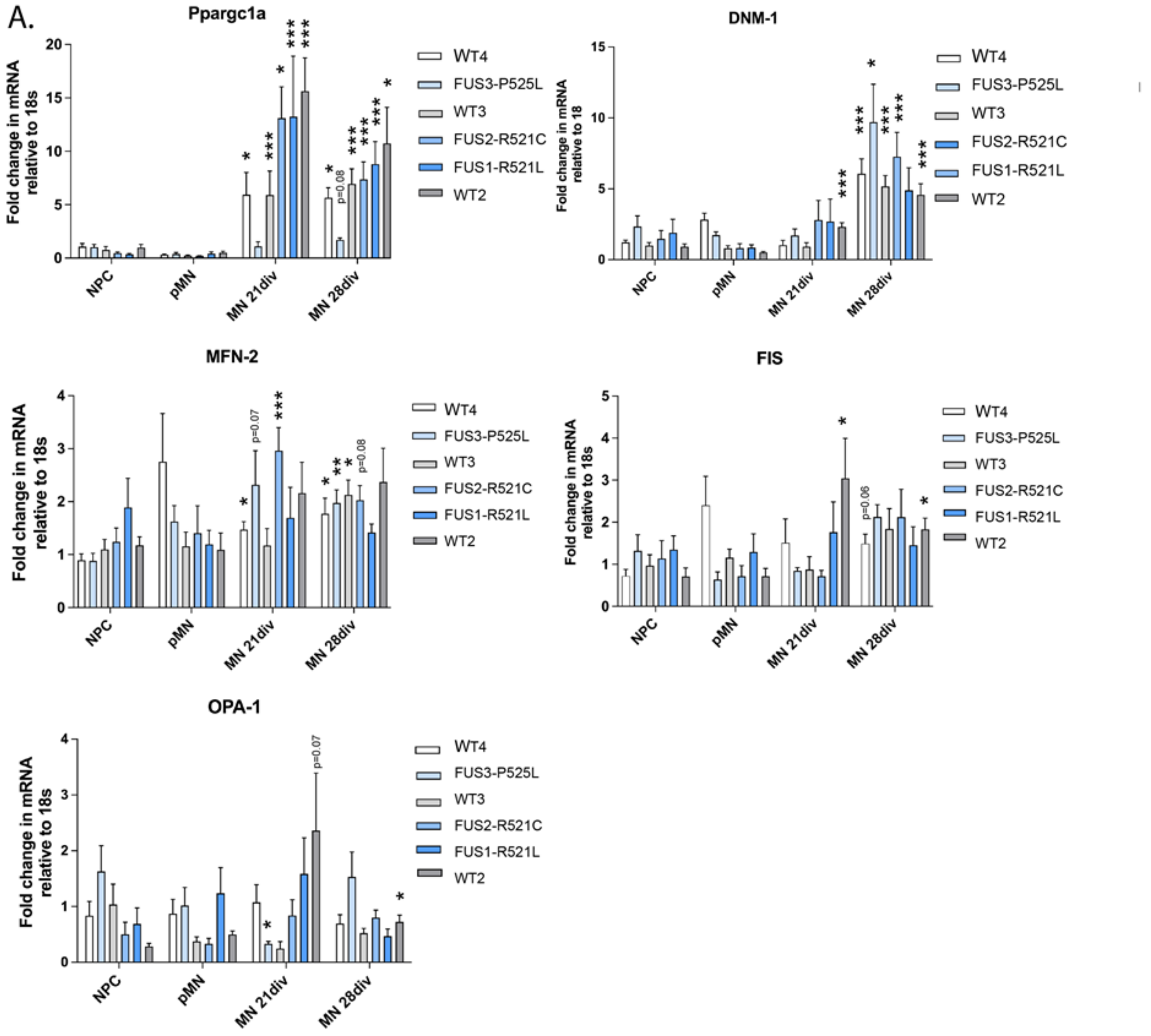


**Supplementary Figure 3. Changes in expression levels of mRNAs important for mitochondrial function during FUS-ALS MN differentiation and maturation. A.**qPCR quantification of relative expression levels in several mRNAs critical for maintenance and operation of the mitochondrial network: PPARGC-1$\alpha$, Dnm1L, MFN-2, OPA1 and FIS1 during MN differentiation and maturation in FUS-ALS lines. Asterisks show significance comparing values between different stages of differentiation within the same line.

Supplementary Figure 4.

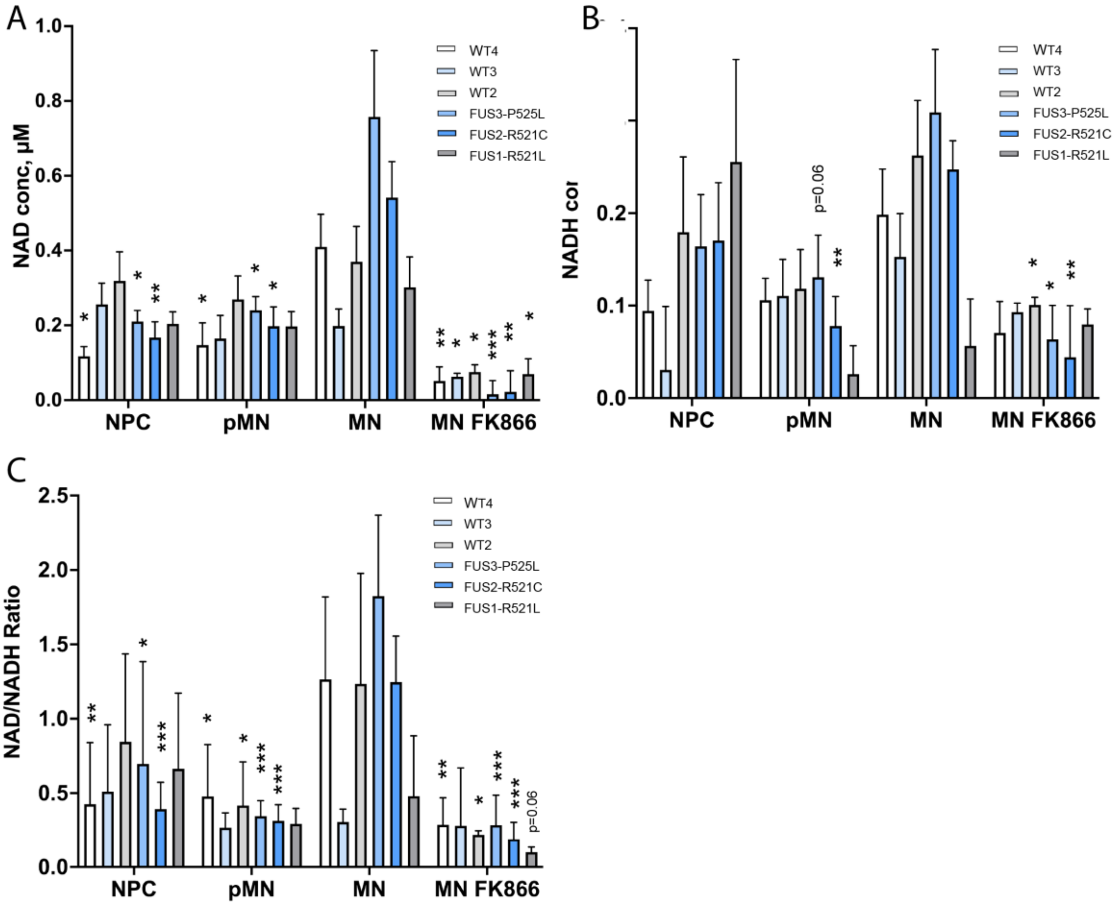


**Supplementary Figure 4. NAD+/NADH levels and redox ratio during MN differentiation of FUS-ALS cultured motoneurons. A-C** Measurements of $NAD^+$ **(A)**, NADH **(B)** concentrations and their redox ratios **(C)** across all tested individual control and FUS mutant lines at different stages of MN differentiation. Significances are shown as * between MN stage vs other stages or treatment within the same MN

### Supplementary Figure 5.

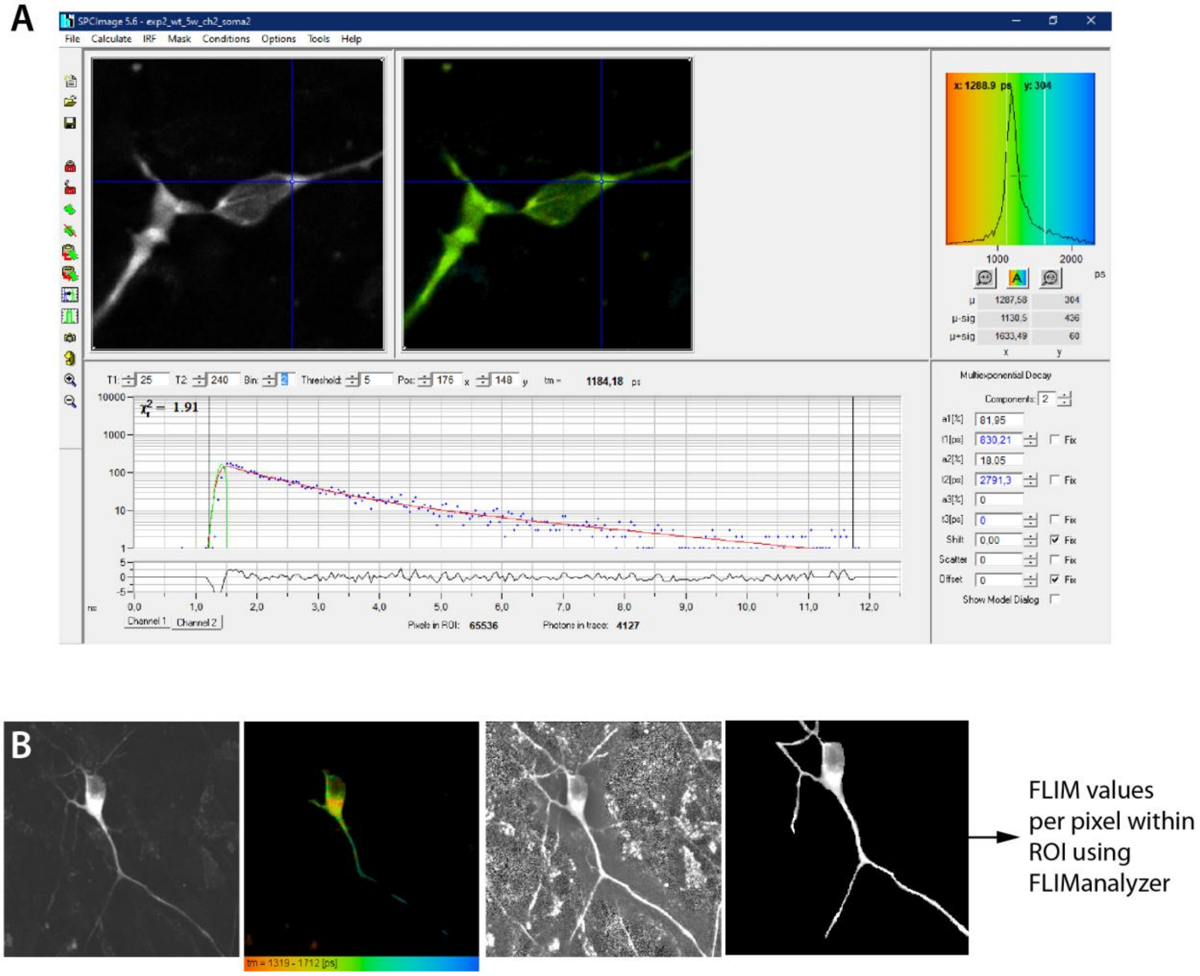


**Supplementary Figure 5. Workflow of FLIM values extraction from single pixel and individual compartments ROI. A.** Snapshot of SPCImage software window with an example curve fitting for single pixel FLIM measurements. Modelling is based on two components incomplete model fitting. **B.** Fluorescence donor (CFP) intensity signal, color coded FLIM values per pixel, and contrast-guided selection of ROI for data extraction to exclude nuclei and background noise measurements.

Supplementary Figure 6.

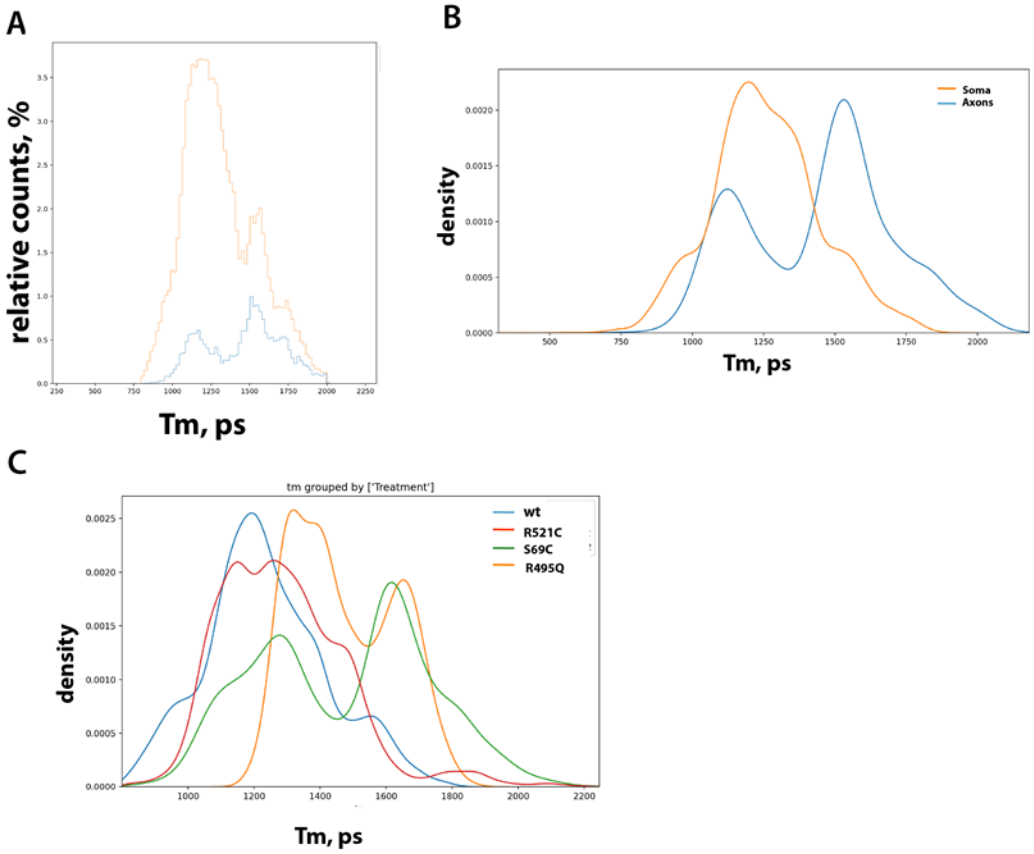


**Supplementary Figure 6. Comparison of individual pixel Tm values distributions in FLIM measurements between somatic and axonal compartments.**

**A.** Frequency distribution histogram **(A)** and normalized KDE distribution plot **(B-C)** comparing Tm values of somatic and axonal compartments **(B),** as well as comparing control (wt1) and mutant FUS MN.

Supplementary Figure 7.

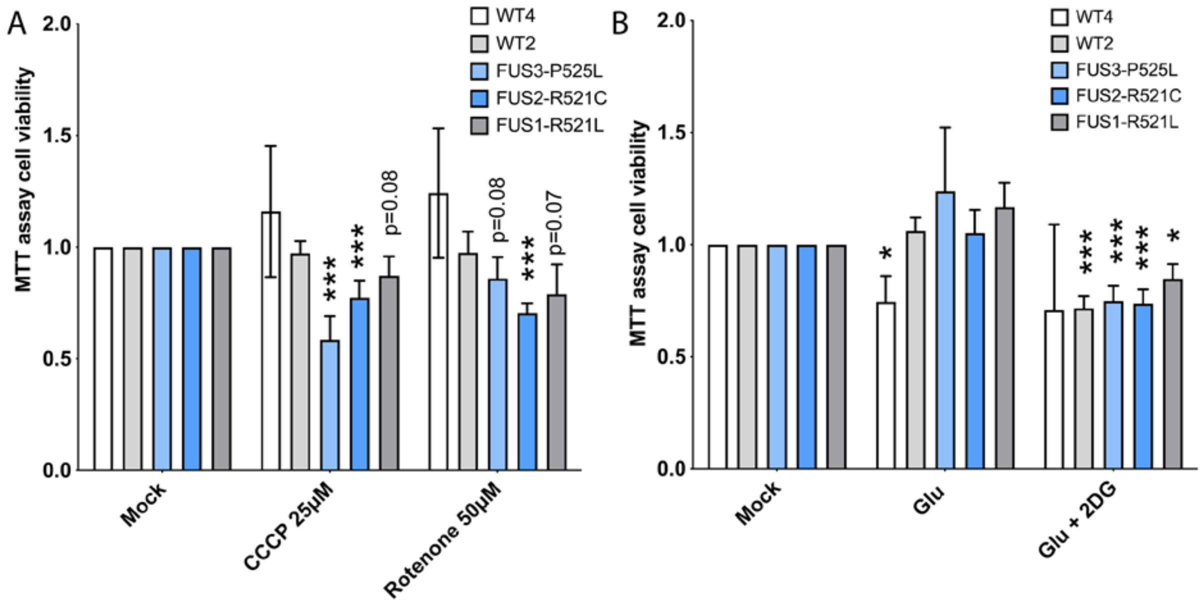


**Supplementary Figure 7. FUS ALS motoneurons have reduced viability upon additional mitochondrial stress. A.** Bar graphs comparing cell survival of individual control and FUS ALS mutant MNs using MTT cell viability assay treated with mitochondrial inhibitors CCCP and Rotenone. **B.** Bar graphs comparing cell survival of individual control and FUS ALS mutant MNs using MTT cell viability assay after glutamate stimulation or blocking glycolysis with 2-deoxy-d-glucose (2DG).
